# Integrated Phenomics and Genomics reveals genetic loci associated with inflorescence growth in *Brassica napus*

**DOI:** 10.1101/2023.03.31.535149

**Authors:** Kevin Williams, Jo Hepworth, Bethany S Nichols, Fiona Corke, Hugh Woolfenden, Pirita Paajanen, Burkhard Steuernagel, Lars Østergaard, Richard J Morris, John H Doonan, Rachel Wells

## Abstract

A fundamental challenge to the production of climate-resilient crops is how to measure dynamic yield-relevant responses to the environment, such as growth rate, at a scale which informs mechanistic understanding and accelerates breeding. The timing, duration and architectural characteristics of inflorescence growth are crucial for optimising crop productivity and have been targets of selection during domestication. We report a robust and versatile procedure for computationally assessing environmentally-responsive flowering dynamics. In the oilseed crop, *Brassica napus,* there is wide variation in flowering response to winter cold (vernalization). We subjected a diverse set of *B. napus* accessions to different vernalization temperatures and monitored shoot responses using automated image acquisition. We developed methods to computationally infer multiple aspects of flowering from this dynamic data, enabling characterisation of speed, duration and peaks of inflorescence development across different crop types. We input these multiple traits to genome- and transcriptome-wide association studies, and identified potentially causative variation in *a priori* phenology genes (including *EARLY FLOWERING3)* for known traits and in uncharacterised genes for computed traits. These results could be used in marker assisted breeding to design new ideotypes for improved yield and better adaptation to changing climatic conditions.

## Introduction

Most plants have evolved to sense environmental cues to coordinate diverse aspects of their development. For example, many species adapted to temperate climates adjust the timing of the initiation and duration of flowering in spring by responding to winter cold, a process called vernalization (Xu and Chong, 2018). Many key crops display this behaviour, with crop types and varieties within species domesticated to different climate zones and agronomic practices. Ensuring crops are locally adapted will be a key challenge to plant breeding as climate change takes effect, for which knowledge of the genetic variants that underlie yield-controlling traits, and their response to temperature, will be critical (Zhu et al., 2022). While model systems have revealed a mechanistic framework (Duncan et al., 2015; Antoniou-Kourounioti et al., 2018), the extent to which such knowledge can be transferred to crops remains unclear (Parent and Tardieu, 2012; Zhu et al., 2022).

The globally-grown crop *Brassica napus* includes a wide range of types such as kales, swedes and winter oilseed rape (OSR), which are grown in cold to temperate winters, semiwinter OSR varieties grown in climates with mild winters, and rapid-cycling crops such as spring OSR, which is grown in short-rotation spring-to-summer conditions. These crop types differ in their flowering time in response to vernalization, a clearly demarcated event that lends itself to traditional manual phenotyping by observational scoring (Meier et al., 2009). However, vernalization is also known to have major effects on shoot architecture and yield (Brown et al., 2019; Hepworth et al., 2020). Traits that are not easy to quantify manually, such as duration and peak of flower production, form critical components for final yield (Kirkegaard et al., 2018). The accurate description of inflorescence dynamics across populations of diverse genotypes is thus valuable, but technically challenging. In agricultural and breeding settings, photogrammetry approaches using drones are under active development, with reasonable predictions of yield (Fang et al., 2016; Gong et al., 2018; Wan et al., 2018). Combined manual assessment and drone imaging has been used to dissect the role of vernalization in floral meristem initiation, including by warming plots (O’Neill et al., 2019). However, although drone-based methods have the potential to be scaled to analyse large populations directly in the field, their spatial resolution is limited and their perspective is restricted to only top view. Moreover, while temperature treatments can be investigated in the field (O’Neill et al., 2019), this is difficult to do at scale across populations.

Proximal imaging provides much higher spatial resolution and with the development of mechanised phenotyping platforms (reviewed by Yang et al., 2020), there is the potential to scale image acquisition to assess many hundreds of individuals on a regular periodic basis. These automated phenotyping systems provide the opportunity to quantify growth and development across time, otherwise known as longitudinal phenotyping. The time dimension can be critical for understanding complex biological processes and the causative linkage of traits. Its importance is being recognised in crop science; Chen et al. (2014) used automated imaging of barley varieties to demonstrate that the broad-sense heritability for many growth and drought response traits was dynamic and changed with growth stage, depending on the particular trait. We have previously found that many growth and developmental quantitative trait loci (QTL) in wheat exert their effects during relatively restricted periods of the life cycle, and some QTL would not have been identified by single measurements (Camargo et al., 2016). Early growth traits have also been found to be good predictors of subsequent development and final yield QTL in oilseed rape (Knoch et al., 2020; Li et al., 2020). Combined with appropriate environmental control and monitoring, automated phenotyping can assess the effect of early treatments, such as vernalization, on subsequent developmental processes across the entire lifecycle.

Genome-wide association studies (GWAS) can locate natural variation within the genome that defines differences in quantitative traits across diversity populations, both to identify variation of value to breeders and to advance knowledge of the genetic basis of diversity (Alseekh et al., 2021). GWAS techniques have been used to identify candidate loci for many traits in *Brassica napus*, including flowering time, plant height and yield (for example; Schiessl et al., 2015; Raman et al., 2016; Li et al., 2018; Wu et al., 2019). The diversity association approach can also be extended to transcriptional variation (Harper et al., 2012; Havlickova et al., 2018; Woodhouse et al., 2021). This ‘Associative Transcriptomics’ (AT) or ‘TWAS’ approach directly links variation in gene expression to the trait and has successfully been used to identify mechanistic variation controlling seed size and crop yield in *B. napus* (Miller et al., 2019). GWAS generates chromosomal regions of interest, within which a causative variant may be found, but does not identify the molecular mechanism concerned. AT does not indicate the causative genetic loci, but does suggest the gene networks involved, and can show high power for variation discovery in smaller populations than are commonly used for GWAS (Harper et al., 2012; Havlickova et al., 2018). A number of studies in OSR combining both techniques have exploited these complementary features to map from genomic variant to molecular mechanism, including the control of flower timing using manual measurement of anthesis (Havlickova et al., 2018; Miller et al., 2019; Raman et al., 2019). Many of the major loci identified by association studies are known flowering regulators in the model plant *Arabidopsis thaliana*, including orthologues of the major floral integrators *FLOWERING LOCUS C* (*FLC)* and *FT* (Raman et al., 2019; Schiessl et al., 2019; Wu et al., 2019; Song et al., 2020). Consistent with the idea that flowering time is critical to agronomic success, allelic variation associated with *FT* paralogs affects not only flowering time *per se*, but also a range of yield traits (Raman et al., 2019).

In this report, we examine how different vernalization temperatures affect not only the initiation of flowering but also subsequent inflorescence development. We use this to explore how the wide range of crop types that represent *B. napus* respond to vernalization in both phenological and morphological characteristics. Finally, we illustrate the potential of high-resolution time-resolved phenotyping combined with GWAS and association transcriptomics to reveal both known and novel candidate genes involved in the variation of known and novel flowering and development traits in *B. napus*.

## Results

### Development of a high-throughput imaging pipeline to estimate the effect of different vernalization conditions on flowering traits

To quantify the effects of vernalization on flowering in *Brassica napus*, a set of accessions was chosen from the Renewable Industrial Products from Rapeseed (RIPR) genetic diversity panel, to represent the breadth of *Brassica napus* diversity while being weighted towards winter oilseed rape, which is defined in part by its requirement for vernalization (Havlickova et al., 2018).

Young plants were subjected to two different temperature regimes in controlled environment rooms, chosen to be climatologically relevant to the UK and north-west Atlantic seaboard of Europe. The plants were then transferred to a mechanised platform in a greenhouse where subsequent growth and development were photographed periodically over eight weeks, resulting in 13888 images from each of three angles.

In order to efficiently process this large dataset, we developed computational procedures to extract features from the time-stamped images. We developed robust and simple algorithms to segment plant images from their background, and subsequently identify and quantify the flowering regions, to provide quantitative data on flowering related traits, typically illustrated by image processing examples in Figure 1 (see Materials and Methods and Supplementary Methods).

**Figure 1.**
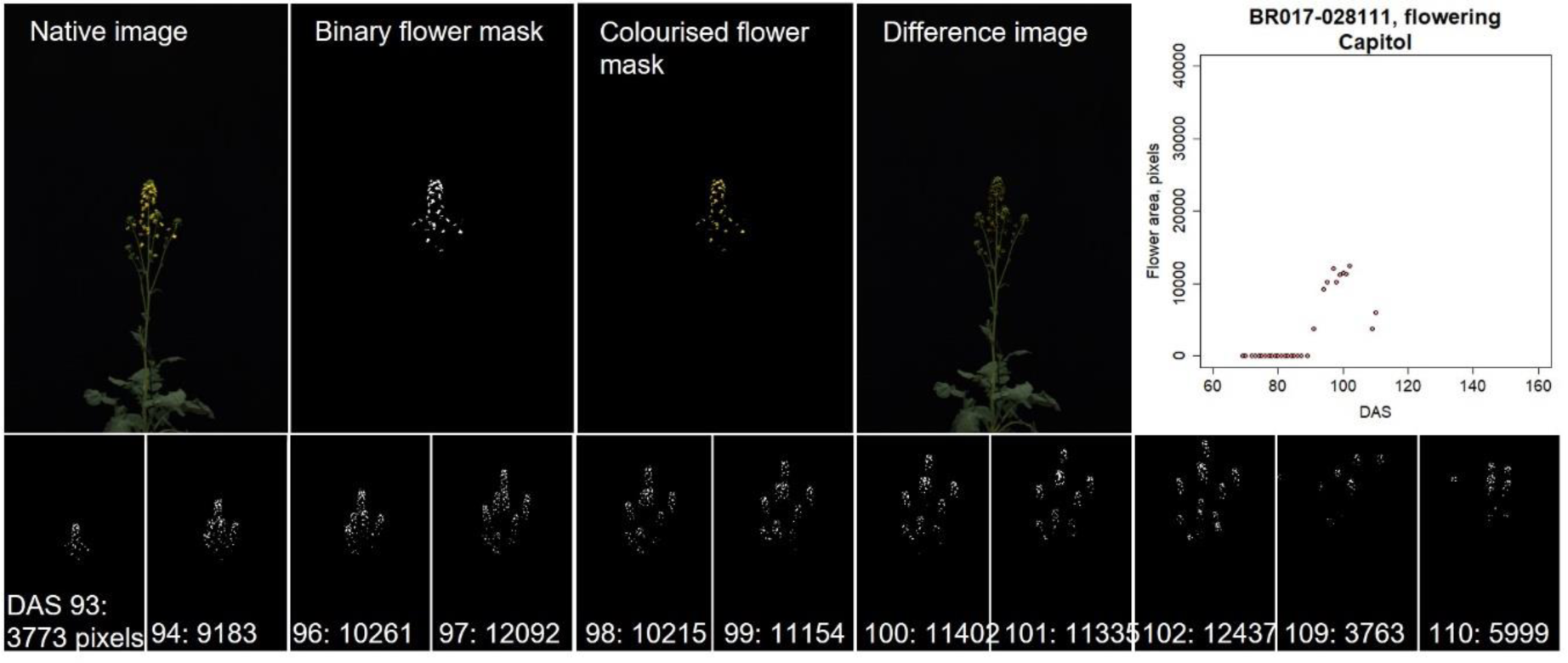
Workflow for segmentation of images for data analysis. Example of segmentation of a single plant (accession “Capitol”) treated with 5°C vernalization, showing an original image, the image after masking has been applied, a series of images over time, and the extracted data plotted over time.

Digital analysis lends itself to extracting traits that are not easy to manually measure on a large scale, such as post initiation floral behaviour. Flowering curves were derived from dynamic image data (Figure 1, 2A and Supplemental Figure S1), and numerical descriptors were calculated for aspects of these curves to explore the nature of flower production over the plants’ lifetime. Flowering curves for each plant were parameterised independently and described nine features, which we have defined as F01 to F09 (Figure 2B and Table 1).

**Figure 2.**
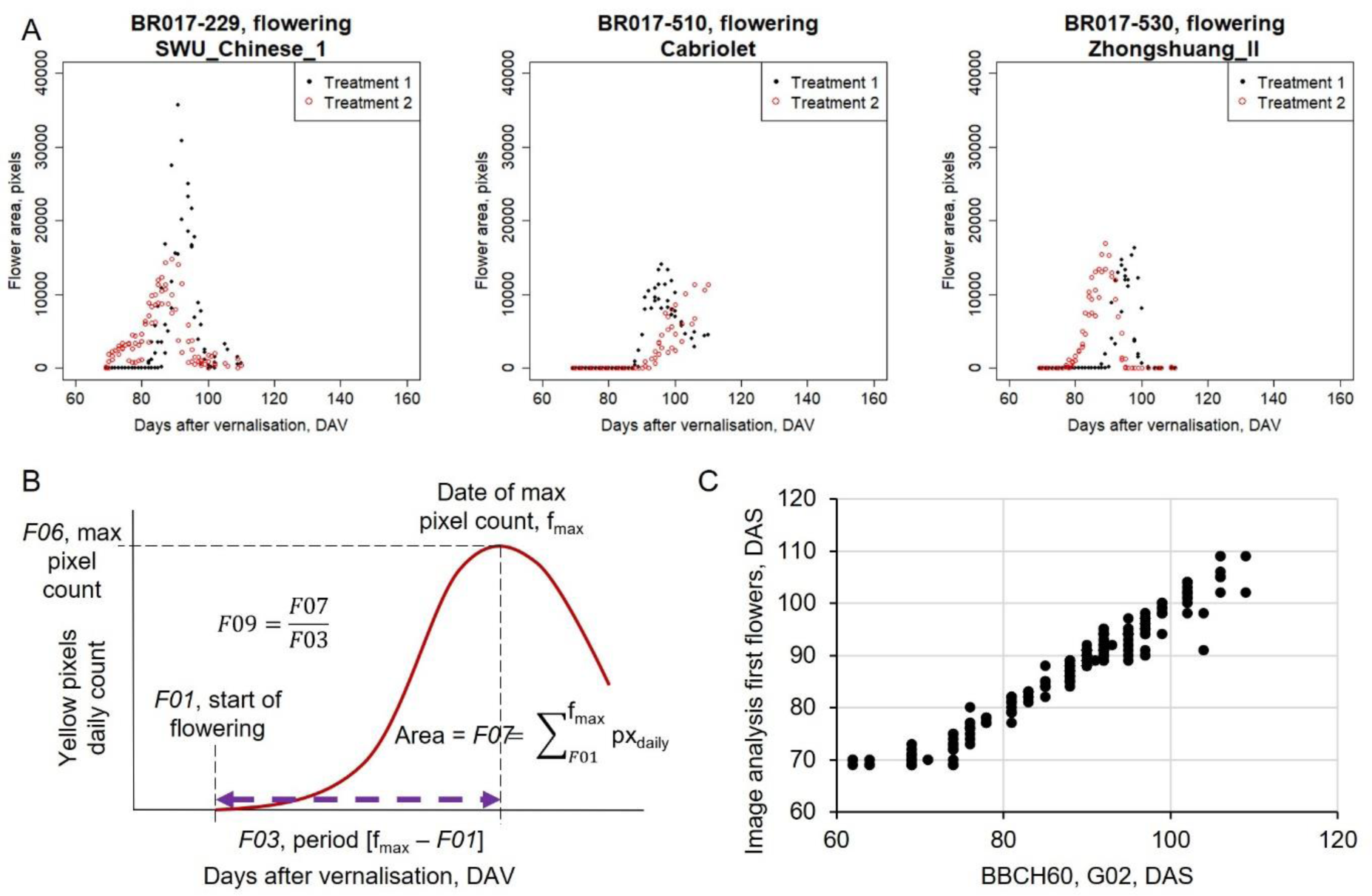
Computational identification of flowering characteristics. A. Flower area by yellow pixel number from side view images, effect of vernalization on three genotypes (combining three replicates per treatment; black = Treatment 1, 5°C vernalization; red = Treatment 2, 10°C vernalization). B. An idealised plot of flowering progression from side view images, showing extracted phenotypes. C, Comparison of first flower phenotype extracted from side view images versus manual scoring. Outliers in the early automated analysis can be seen where flowering commenced before imaging (earlier than 69 days after sowing).

**Table 1.** Features extracted from high-throughput phenotyping by dynamic image capture and comparative ground truth measurement. G## coded traits indicate “ground truth” manually scored flowering traits. F## codes indicate traits derived from the flower pixel analysis, while R## codes derive from the raceme analysis. Px = pixels.

| Description of trait | Code | Unit | Computational name |
| --- | --- | --- | --- |
| Manual observation of DAS to the appearance of visible flower buds vertically from the apex | G01 | Days | BBCH51 |
| Manual observation of DAS to the appearance of open flowers | G02 | Days | BBCH60 |
| Time from manual BBCH51 (G01) to manual BBCH60 (G02) observations | G03 | Days | Ground_BBCH60-BBCH51 |
| Number of days between sowing and appearance of first flower pixels (BBCH60) | F01 | Days | first_flowers_DAS |
| Number of days between sowing and the date when most flower pixels were counted | F02 | Days | peak_flowers_DAS |
| Number of days between the first flower and peak flowering | F03 | Days | flowering_days_to_peak |
| Number of days from the first to last flowers being recorded | F04 | Days | total_flowering_days |
| Had the flowering already finished on the day that the plant was removed from the LT system? | F05 | Yes/no | flowering_finished |
| Number of flower pixels seen on date of peak flowering | F06 | Px | pixel.day_peak |
| Aggregate of all flower pixels (including interpolated data) from the date of first flowering to the peak | F07 | Px | pixels.days_to_peak |
| Aggregate of flower pixels across all days (true images plus interpolated data) | F08 | Px | total_pixel.days |
| Arithmetic mean of pixels per day from first flower to peak flowering | F09 | Px | mean_pixels.day_to_peak |
| Time from manual BBCH51 (G01) to automated BBCH60 (F01) observations | F10 | Days | Auto_BBCH60-BBCH51 |
| Onset of raceme extension (datestamp when first observed poking above canopy) | R01 | Days | raceme_start |
| Mean rate of extension through growth phase up to 90% of the maximum height | R02 | Px.day<sup>-1</sup> | raceme_extension_rate_90 |
| Maximum rate of raceme extension | R03 | Px.day<sup>-1</sup> | max_extension_rate |
| DAS (or DAV) at which R03 occurs | R04 | Days | max_rate_DAS |
| Number of days of raceme growth to reach 90% of max height | R05 | Days | raceme_growth_period_90 |
| Number of days of raceme growth to reach 50% of max height | R06 | Days | raceme_growth_period_50 |
| Number of days of raceme growth to reach 75% of max height | R07 | Days | raceme_growth_period_75 |
| Mean rate of extension through growth phase up to 50% of the maximum height | R08 | Px.day<sup>-1</sup> | raceme_extension_rate_50 |
| Mean rate of extension through growth phase up to 75% of the maximum height | R09 | Px.day<sup>-1</sup> | raceme_extension_rate_75 |
| Maximum plant height | R10 | Px | max_height |
| DAS at which R10 occurs | R11 | Days | max_height_DAS |
| Period between start of raceme extension and peak height | R12 | Days | extension_days_to_max |
| Flag to indicate raceme collapse | R13 | 0/1 | raceme_collapse |
| Time from manual BBCH51 (G01) observations to automated raceme start (R01) | R14 | Days | BBCH51_to_raceme_start |
| Flag to indicate elongation started during vernalization | R15 | 0/1 | elongation_during_vernalization |

We also manually scored the commonly-used phenology traits “BBCH51” (floral buds visible from above) and “BBCH60” (first flower opening) (Meier et al., 2009) as “ground truth” data to monitor experimental progress and test our digital analysis. Floral initiation data (G02) were compared directly to the first appearance of yellow flower pixels (F01) in each dataset. Strong linear correlation was seen between manual BBCH60 scoring and the onset of flowering as calculated through side view image analysis [Days After Sowing (DAS): *y* = 0.94*x* + 5.1 (*R*^2^ = 0.95), where (70 < *x* < 110), suggesting a typical deviation of ±1 day between techniques] (Figure 2C). Similarly, a strong correlation was confirmed when comparing side to top view images [DAS: *y* = 1.01*x* - 0.4 (*R*^2^ = 0.98)] (Supplemental Figure S2). Verification of the analysis for floral initiation promotes confidence that our subsequent observations are accurate.

While analysing floral dynamics, it became apparent that there was variation between plants in the rate of primary stem (raceme) elongation. Therefore, we developed a specific sub-routine to quantify elongation and extract features describing raceme timing, speed and character traits (Supplemental Figure S3). Further features were extracted from the manual ‘ground-truth’ features (G01-G02), manual data-cleaning (F05, R13, R15) and from comparing these manual traits with each other and the automated feature extraction (G03-F10, R14). In all, our manual phenotyping, flower detection and raceme tracking resulted in a total of twenty-nine phenotypes describing the dynamic development of the inflorescence, with two pairs of traits directly duplicated between automation and human observation (G02-F01, G03-F10). The results for each plant are shown in Supplemental Tables S1-4.

### High Throughput Imaging reveals differences in both phenological and morphological responses to vernalization temperature between crop types

Having developed the phenotyping capacity, we investigated how these phenotypes varied across the population. In both the manual and automated phenotyping, expected responses in floral initiation to vernalization across crop types were confirmed, with winter OSR and plants grown for pre-flowering harvest (leafy vegetables, winter fodder and swedes) generally flowering later than spring and semiwinter OSR (G01, F01; Figure 3, S4). On average, the winter OSR lines show a slight acceleration in flowering in response to vernalization at 5°C compared to 10°C, as expected from their vernalization requirement, whereas spring and semiwinter OSR lines are strongly delayed at 5°C.

**Figure 3.**
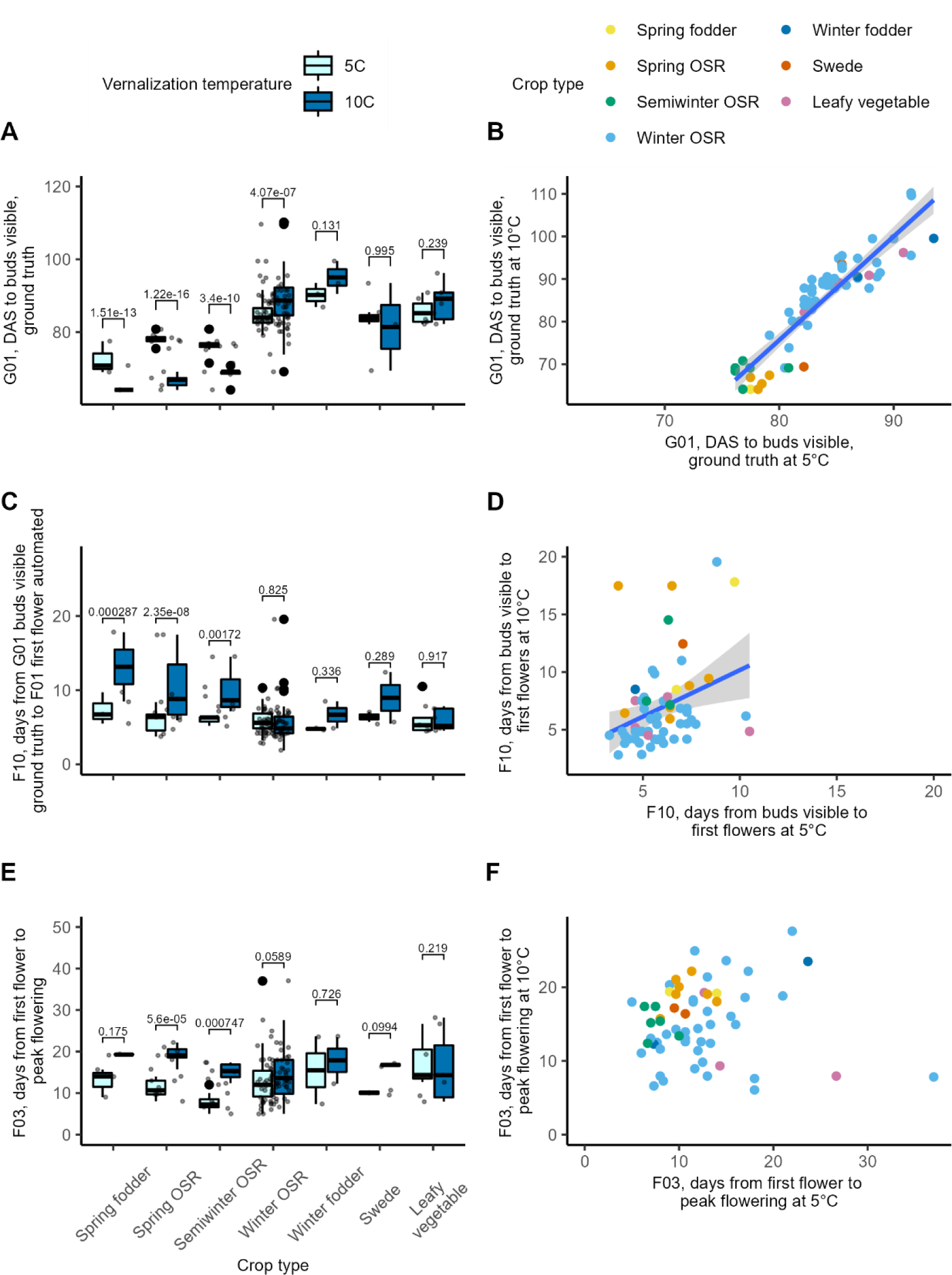
Phenological responses to vernalization temperature across different crop types. Comparison of responses to different vernalization temperatures for: A-B) G01 – days from sowing to buds visible ‘BBCH51’ stage; C-D) F10, days from buds visible (manual) to first flower opening by automated analysis; E-F) F03 – days from automated first flower scoring to peak yellow pixels. Panels A, C, E show boxplots by crop type and vernalization treatment, p-values are Tukey-corrected pairwise contrasts per crop type for response to vernalization temperature (see Methods). Panels B, D, F show scatter plots with trendline showing linear regression; B) R^2^=0.856, p-value= < 2.2×10^−16^; D) R^2^=0.105, p-value=0.00553; F) trend not shown, p-value: 0.892. Points on all graphs show genotype estimated marginal means per accession. N varies per treatment and trait, but maxima N are: Spring fodder = 3, spring OSR = 8, semiwinter OSR = 8, winter OSR = 42, winter fodder = 2, swede = 2, leafy vegetable = 6.

There was a very tight correlation between buds visible (G01) response to temperature across the different lines, suggesting that genotypes that flower later in response to one temperature, also flower relatively late in response to another – i.e. winter OSRs are always slower than springs (Pearson’s correlation = 0.927, p-value < 2.2×10^−16^; Figure 3B). This trend continues into later absolute-timing traits, such as days to first flower (G02, Pearson’s = 0.873, p-value < 2.2×10^−16^, Figure S4); into the automated analysis of equivalent traits (G01 ≈ R01 bolting start, for which Pearson’s = 0.884, p-value < 2.2×10^−16^, G02 = F01 first flower F01, for which Pearson’s = 0.877, p-value < 2.2×10^−16^, Figure S4); and into the new automated traits that also define absolute timing: days to maximum extension rate (R04, Pearson’s = 0.874, p < 2.2×10^−16^), days to peak flowers (F02, Pearson’s = 0.803, p = 2.14×10^−15^) and days to maximum height (R11, Pearson’s = 0.844, p < 2.2×10^−16^; Figure S5).

There is more variation in the timing between different developmental stages. For example, the time between buds visible and first flowers for each accession is much more variable in response to temperature than either absolute timing trait (F10, Pearson’s = 0.346, p = 0.00553; Figure 3C-D, Figure S4). While late-flowering crops have no temperature response on average in bolting-to-flowering time, semiwinters and springs are significantly delayed in response to 10°C treatment, suggesting that warmer winters accelerate bolting more than flower opening. A similar pattern is seen for days from buds visible to start of raceme extension (R14; G01 to R01; Figure S5) and particularly days from first flower opening to peak (F03; F01 to F02, Figure 3E-F), with inter-genotype variability having increased even further in this later trait.

The rates of raceme extension (R02, R03, R08 and R08) are significantly higher in response to 5°C vernalization than 10°C across almost all crop types, but vary widely within each crop (Figure 4A, Figure S6). Conversely, the duration of extension varies much less between treatments (R05, R06, R07; Figure 4C, Figure S6), with the total period of extension (R12) highly similar across lines and treatments; semiwinter OSR and winter fodder are the only groups showing much response (Figure S6). The faster rates of extension resulted in taller crops post 5°C than post 10°C, regardless of type, with winter crops generally taller than springs (Figure 4E).

**Figure 4.**
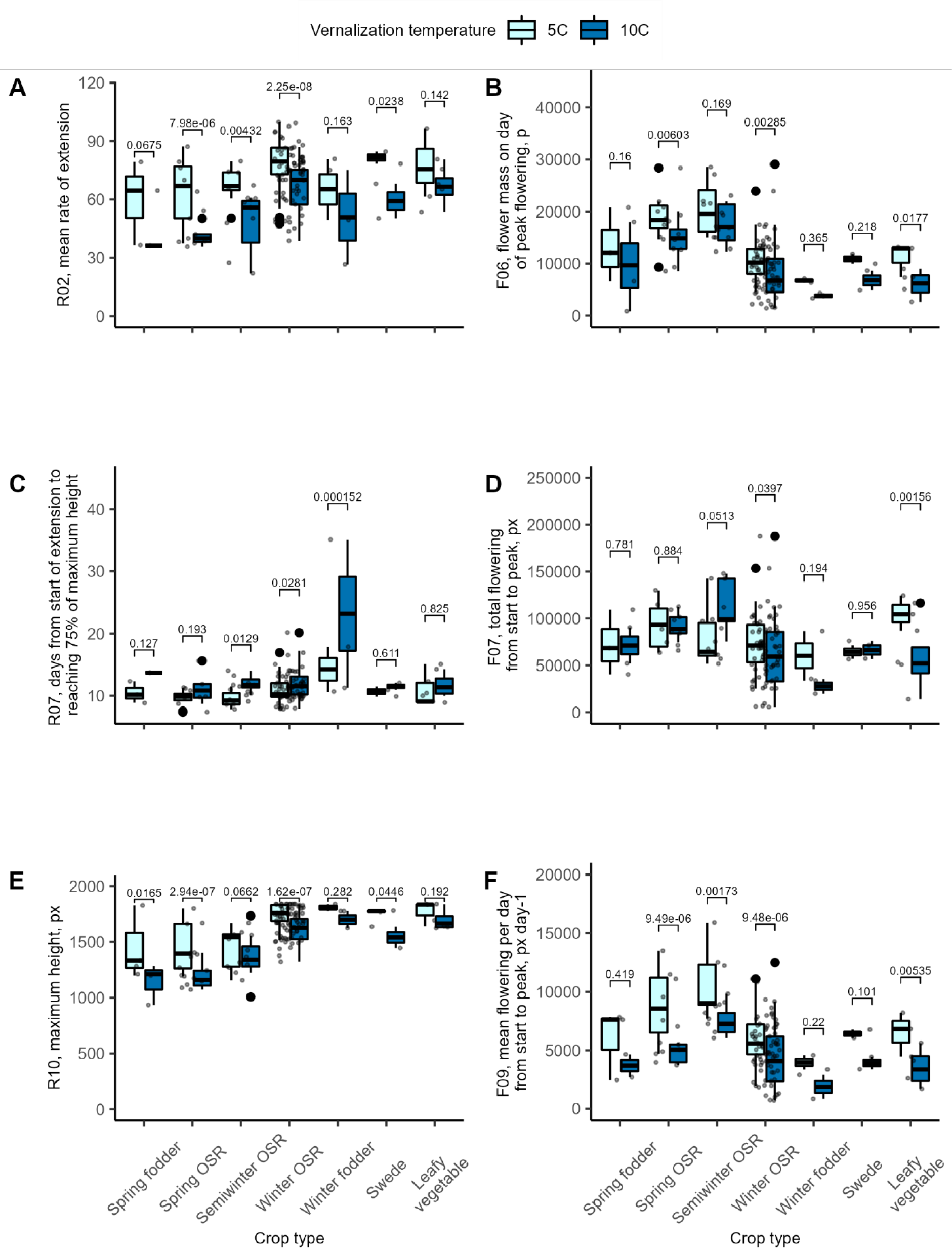
Morphological responses to vernalization temperature across different crop types. Comparison of responses to different vernalization temperatures by crop type for; A) R02, mean rate of raceme extension per day; B) F06, flower area at peak, in pixels; C) R07, days from the start of raceme extension to reaching 75% of the maximum height; D) F07, total flower area accumulated from start of flowering to peak flowering, in pixels; E) R10, maximum height in pixels (px); F) F09, mean flower area per day from start of flowering to peak flowering, in pixels per day. Boxplots by crop type and vernalization treatment, p-values are Tukey-corrected pairwise contrasts (see Methods), points represent genotype estimated marginal means. N varies per treatment and trait, but maxima N are: Spring fodder = 3, spring OSR = 8, semiwinter OSR = 8, winter OSR = 42, winter fodder = 2, swede = 2, leafy vegetable = 6.

Vernalization at 5°C results in faster raceme extension, but within a more compressed time period: a similar process seems true of flower production (Figure 4B, D, F). Although the flowering time from start to peak is much shorter in spring and semiwinter OSR after 5°C treatment (F03; Figure 3E), the mean flowering area per day (F09; Figure 4F) is increased across all OSR by 5°C treatment. However, the greater duration of flowering in response to 10°C vernalization in spring OSR and swedes is such that total flowering over this period (F07; Figure 4D) is very similar between treatments, while semiwinters flower more after 10°C. As flowering was not completely finished at the termination of the experiment, trait F07 (total flower area to peak) is the most likely trait of those investigated here to correspond to yield for the OSR varieties (Kirkegaard et al., 2018; Siles et al., 2021). While 5°C vernalization generally promotes flower production in winter crops as expected, it does not much affect the total flower area for springs, while the ‘facultative responding’ semiwinters show a marked improvement in response to warmer vernalization temperatures – and the highest flower area of all on average.

We investigated how different traits relate to each other within temperature categories. Absolute timing traits are all closely related (G01, G02, F01, F02, R01, R04, R11: Supplemental Figure S7). Traits at 10°C covary more closely and significantly than at 5°C, though the patterns are largely similar. We calculated values for temperature responses both by subtraction (10-5) and division (10v5; for this comparison, only post-vernalization times were compared). Comparing traits by proportion (10v5) rather than subtraction (10-5) creates many more significant correlations, suggesting that this is more biologically meaningful (Supplemental Figure S7). Traits describing flower area (F06, F07 and F09) positively covary with each other but are negatively related to timing traits – probably reflecting the higher flower area of the spring and semiwinter varieties. This trend may explain why many covariances are stronger after 10°C vernalization than 5°C, as warmer conditions increased the variation between crop types. Within raceme traits, extension duration traits (R05, R06, R07) are usually negatively correlated with rate (R02, R03, R08, R09), suggesting an intrinsic trade-off between the speed and the duration of growth.

### GWAS identifies candidate chromosomal regions

To investigate the genetic bases of these traits and their environmental response, we employed a genome wide-association study approach. A limitation in large-scale automated phenotyping producing multiple traits is the complexity of running, model fitting and analysing so many resulting GWA studies. To expedite this, we developed an analysis pipeline (GAGA: Nichols et al., in prep.) to efficiently process the resulting 98 traits for both GWAS and Associative Transcriptomics. This pipeline outputs a list of significantly associated single nucleotide polymorphism (SNP) and gene expression markers (GEMs), saving analysis time.

Significantly associated markers are identified for several traits, some of which are close to promising targets, particularly for the 10°C treatment, validating the approach (Figure 5, Table S5). Targets with clear association peaks include F03 at 10°C, where the most significant associated region on C09 (marker Bo9g018460.1:276:G) takes in homologues of *ABA REPRESSOR1* and *EARLY FLOWERING5* (*AT5G64750* and *AT5G64813* respectively); genes controlling growth rates, cold-responses and flowering time (Noh et al., 2004; Pandey et al., 2005). Other cases show single significant markers without linkage-dragged peaks but are close to strong candidates, and often are supported in several GWAS models. For example, at 10°C for G01 the marker Bo4g023950.1:823:C is within 1kb of *BnaC04g05100*, orthologue of *At2g44710,* an RNA binding protein that affects *FLOWERING LOCUS C* (*FLC*) levels, a master regulator of flowering time (Zhang et al., 2022). R04 and R11 at 10°C and R04 in the 10v5 treatment share a region of significance on A08 (markers in Cab036002.1, BnaA08g11330D, Cab036009.2, Cab036011.1, and Cab036069.1), which though has no clear regulatory candidates, does show synteny with the Arabidopsis region harbouring *AT4G34400, TARGET OF FLC AND SVP1,* which modulates the floral transition (Richter et al., 2019). For R11 at 10°C, a peak on A03 (Cab015902.1:984:T; Figure 5) is in the gene model next to an *AGAMOUS LIKE15* homologue (*AGL15*, *BnaA03g04490D, AT5G13790*), while an orthologue of *AGL19* is the site of the highest marker for G02 in the 10-5 treatment (Cab035904.1:314:T, *BnaA08g10540D*, *AT4G22950*). The strongest marker for G03 in 10v5 (Cab024343.4:1647:C; Chr. A01) is adjacent to a gene orthologous to *AGL24* (*BnaA01g13920D, AT4G24540*), a key interactor of *SUPPRESSOR OF OVEREXPRESSOR OF CONSTANS1* (*SOC1*; itself also known as *AGL20*). As regulators of the floral transition (Pajoro et al., 2014; Richter et al., 2019), all the *AGLs* are strong candidates for these timing traits.

**Figure 5.**
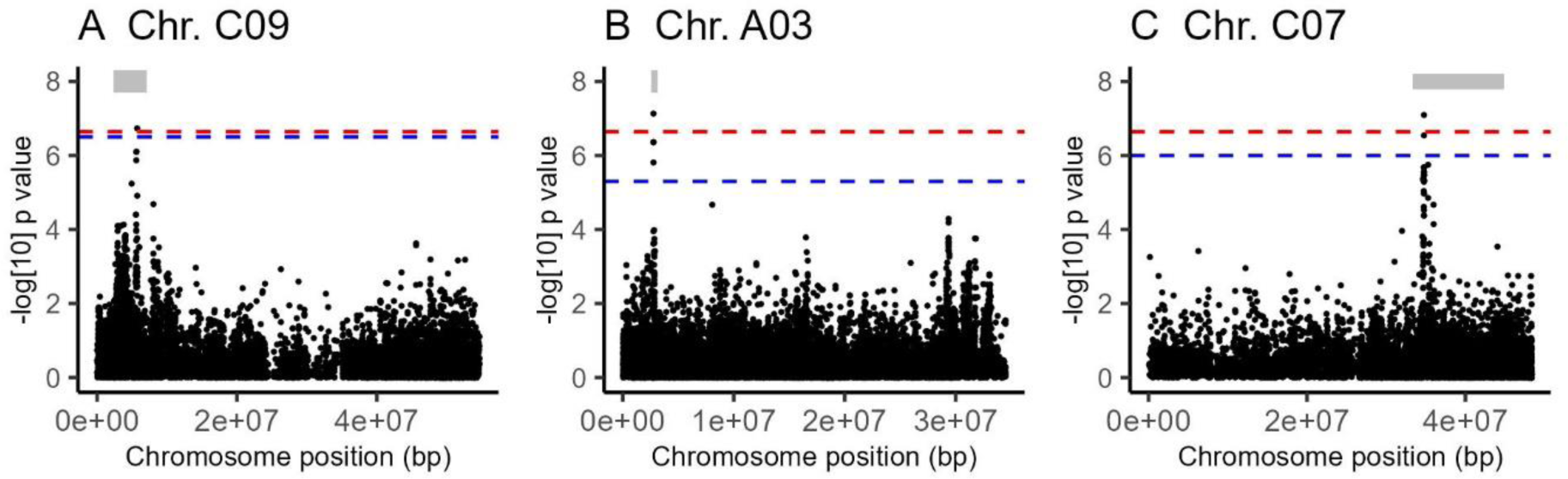
Manhattan plots showing probabilities of a link between Single Nucleotide Polymorphism (SNP) presence versus different traits. Manhattan plots for data with peaks of expression generated by GAPIT via GAGA, for each trait the best-fitting GWAS model (GLM, FarmCPU or BLINK) is shown. Red dashed line marks Bonferroni significance at α = 0.05, blue dashed line shows Benjamini-Hochberg False Discovery Rate (FDR) of α = 0.05, thick grey lines indicate region where linkage disequilibrium (LD) r^2^ >0.2 for the highest marker. A) F03 - days from flowering start to peak yellow pixels after 10°C vernalization, chromosome (Chr.) C09, Blink, LD 2434537-6989280bp. B) R11 - days after sowing to maximum height at 10°C vernalization, Chr. A03, FarmCPU, LD 2605764-3119409. C) F02 - ratio of days to peak flowering after vernalization at 10°C versus 5°C, Chr. C07, FarmCPU, LD 33374075-44866613bp.

For F02 10v5 and 10-5, a clear peak on Chr. C07 (Bo7g089790.1:696:C) includes a number of markers in *BnaC07g27010D,* a gene with homology to *OVULE ABORTION 2* (*OVA2, AT5G49030*), which is required for female reproductive development (Berg et al., 2005). QTLs for flowering time locating to this region has previously been identified in a cross between winter and spring varieties of OSR (Li et al., 2018).

A number of phenotypes did not produce significant signals (Supplemental Table S7). This is probably because of the combination of the small population (69 genotypes were suitable for the analysis, with further losses for some traits due to data curation), population structure, and the highly polygenic nature of several phenotypes.

### AT permits greater power to identify genes characterising quantitative traits

Transcriptome analysis revealed that many traits correlated with expression of a large number of genes, confirming their polygenic character. The phenology traits (G01, G02, F01, R01, and R04) are highly significantly associated with more than 100 genes at both vernalization temperatures (genes significant at Bonferroni-correction p-value < 0.05; Supplemental Table S6).

Some traits still did not produce any significant associations (Supplemental Table S7). At 5°C, almost all these related to different variations of inter-trait timing (F03, F07, R05, R12). Very few traits produced markers for 10-5, but many traits did for 10v5.

There were also overlaps with the GWAS. For F02 (peak flower date) at 5°C the highest GEM and highest SNP marker were both in *BnaA07g09950D*, an *ADP-RIBOSYLATION FACTOR 1A* orthologue, while for the same trait at 10v5, the 18^th^ ranked GEM was in the same gene model (*OVA2)* as the most significant GWAS hit (Table 2).

**Table 2.** Significant a priori Gene Expression Markers.

| Pantranscriptome unigene, Ensembl GI, Name, Arabidopsis GI, Included for exome capture | Traits | FDR corrected p value, bold indicates significant at Bonferroni corrected $\alpha < 0.05$, # indicates ranking. | | | |
| --- | --- | --- | --- | --- | --- |
|  |  | 5°C | 10°C | 10°C-5°C | 10°C/5°C |
| *Cab008705.1* | G01 | 0.00218 | 0.00517 | n.s. | n.s. |
| *BnaA07g09950D* | G02 | 0.00119 | 0.00554 | 0.0171 | 0.0191 |
| *BnaARFA1A.A07* | F01 | 9.72 x10-4 | n.s. | n.s. | n.s. |
| *AT1G23490* | F02 | **9.70 x10-4, #1** | 0.0247 | n.s. | n.s. |
| No | R01 | 0.00270 | n.s. | n.s. | n.s. |
|  | R04 | **2.17 x10-4** | n.s. | n.s. | 0.0304 |
|  | R06 | 0.0138 | n.s. | n.s. | n.s. |
|  | R10 | 0.006045 | n.s. | n.s. | n.s. |
|  | R11 | **0.00511, #4** | n.s. | n.s. | n.s. |
| *Bo7g089790.1* | G02 | n.s. | n.s. | n.s. | 0.0316 |
| *BnaC07g27010D* | F01 | n.s. | n.s. | n.s. | n.s. |
| *BnaOVA2.C07* | F02 | n.s. | n.s. | 0.00526 | 0.00340 |
| AT5G49030 |  |  |  |  |  |
| No |  |  |  |  |  |
| *BnaC02g00490D* | G01 | **1.28 x10<sup>-5</sup>, #1** | **5.45 x10<sup>-6</sup>** | n.s. | **5.41 x10<sup>-6</sup>** |
| *BnaC02g00490D* | G02 | **1.08 x10<sup>-3</sup>** | **1.17 x10<sup>-5</sup>** | **5.73 x10<sup>-6</sup>, #10** | **3.91 x10<sup>-6</sup>** |
| *BnaFLC.C02* | F01 | **1.94 x10<sup>-4</sup>** | **2.04 x10<sup>-5</sup>** | n.s. | **8.02 x10<sup>-6</sup>** |
| *AT5G10140* | F10 | n.s. | 0.0404 | n.s. | n.s. |
| Yes | F02 | 0.037867 | **2.90 x10<sup>-4</sup>, #2** | 0.00707 | 0.00333 |
|  | F06 | **0.00578** | 0.00750 | n.s. | n.s. |
|  | R01 | **1.65 x10<sup>-5</sup>, #3** | **1.85 x10<sup>-5</sup>, #10** | n.s. | **4.73 x10<sup>-5</sup>** |
|  | R04 | **3.30 x10<sup>-5</sup>, #2** | **4.31 x10<sup>-5</sup>** | n.s. | **3.75 x10<sup>-5</sup>** |
|  | R06 | 0.0217 | n.s. | n.s. | n.s. |
|  | R10 | **5.17 x10<sup>-5</sup>** | 0.00289 | n.s. | n.s. |
|  | R11 | 0.00856, #7 | 0.00685 | 0.0260 | 0.0109 |
|  | R14 | n.s. | 0.0349 | n.s. | n.s. |
|  | R15 | n.s. | 0.00243 | n.s. | n.s. |
| *Cab002472.4* | G01 | **1.28 x10<sup>-5</sup>** | **5.45 x10<sup>-6</sup>** | n.s. | **1.76 x10<sup>-5</sup>** |
| *BnaA03g13630D* | G02 | 0.00182 | **1.17 x10<sup>-4</sup>** | **5.73 x10<sup>-6</sup>** | **3.90 x10<sup>-6</sup>** |
| *BnaFLC.A03b* | F01 | 0.000146 | **2.04 x10<sup>-4</sup>** | n.s. | **8.02 x10<sup>-6</sup>** |
| *AT5G10140* | F02 | n.s. | **7.50 x10<sup>-4</sup>** | 0.00692 | 0.00355 |
| Yes | F06 | 0.00562 | 0.0300 | n.s. | n.s. |
|  | R01 | **1.63 x10<sup>-5</sup>** | **3.34 x10<sup>-4</sup>** | n.s. | **2.28 x10<sup>-4</sup>** |
|  | R04 | **3.30 x10<sup>-5</sup>** | **6.81 x10<sup>-5</sup>** | n.s. | **2.41 x10<sup>-5</sup>** |
|  | R06 | **0.00433, #2** | 0.005288034 | n.s. | n.s. |
|  | R10 | **1.80 x10<sup>-5</sup>** | **4.41 x10<sup>-5</sup>** | n.s. | n.s. |
|  | R11 | 0.0221 | **0.00243, #10** | **0.00296, #3** | **4.33 x10<sup>-4</sup>, #4** |
|  | R14 | n.s. | n.s. | n.s. | n.s. |
|  | R15 | n.s. | 0.00860 | n.s. | n.s. |
| *Bo4g024850.1* | G01 | 9.24 x10<sup>-4</sup> | **5.94 x10<sup>-5</sup>** | n.s. | **5.41 x10<sup>-6</sup>** |
| *BnaC04g53290D* | G02 | **1.61 x10<sup>-4</sup>** | 7.36 x10<sup>-4</sup> | 0.00177 | **4.55 x10<sup>-5</sup>** |
| *BnaSOC1.C04* | F01 | **1.51 x10<sup>-4</sup>** | **3.33 x10<sup>-5</sup>** | n.s. | **8.02 x10<sup>-6</sup>** |
| *AT2G45660* | R01 | **2.56 x10<sup>-4</sup>** | **1.98 x10<sup>-4</sup>** | n.s. | 0.00108 |
| Yes | R02 | 0.0442 | n.s. | n.s. | n.s. |
|  | R03 | 0.0451 | n.s. | n.s. | n.s. |
|  | R04 | 0.00117 | 0.00141 | n.s. | 0.00320 |
|  | R09 | 0.0110 | n.s. | n.s. | n.s. |
|  | R10 | 0.00148 | 0.00273 | n.s. | n.s. |
|  | R11 | n.s. | 0.0223 | n.s. | 0.0305 |
|  | R15 | n.s. | 0.0146 | n.s. | n.s. |
| *Bo2g096070.1* | G01 | **1.28 x10<sup>-5</sup>** | **6.31 x10<sup>-6</sup>** | n.s. | **5.41 x10<sup>-6</sup>** |
| *BnaC02g23180D* | G02 | 0.00485 | **1.17 x10<sup>-5</sup>, #1** | **5.73 x10<sup>-6</sup>, #3** | **3.91 x10<sup>-6</sup>, #2** |
| *BnaGA3OX2.C02* | F01 | 0.00133 | **2.04 x10<sup>-5</sup>, #6** | n.s. | **8.02 x10<sup>-6</sup>, #6** |
| *AT1G80340* | F02 | n.s. | **4.26 x10<sup>-4</sup>, #7** | 0.0278 | 0.0112 |
| No | F06 | **0.00156, #4** | 0.00762 | n.s. | n.s. |
|  | R01 | **2.08 x10<sup>-4</sup>** | **1.85 x10<sup>-5</sup>, #5** | n.s. | **2.26 x10<sup>-5</sup>, #6** |
|  | R04 | 0.00112 | **3.02 x10<sup>-5</sup>, #4** | n.s. | **2.41 x10<sup>-5</sup>, #5** |
|  | R09 | n.s. | 0.0169 | n.s. | n.s. |
|  | R10 | 0.00124 | 0.00116 | n.s. | n.s. |
|  | R11 | 0.0210 | 0.00323 | 0.0171 | 0.00483 |
|  | R14 | n.s. | 0.0450 | n.s. | n.s. |
|  | R15 | 0.00968 | 0.0188 | n.s. | n.s. |
| *Cab032577.1* | G01 | **1.08 x10<sup>-4</sup>** | **5.45 x10<sup>-6</sup>** | n.s. | 0.00466 |
| *BnaA02g06490D* | G02 | **4.12 x10<sup>-5</sup>** | 4.69 x10<sup>-4</sup> | 8.27 x10<sup>-4</sup> | 4.17 x10<sup>-5</sup> |
| *BnaTOE2.A02* | F01 | **2.78 x10<sup>-5</sup>** | 0.00554 | n.s. | 0.00463 |
| *AT5G60120* | F02 | n.s. | 0.0392 | n.s. | n.s. |
| Yes | R01 | **9.53 x10<sup>-5</sup>** | 0.00528 | n.s. | 0.00665 |
|  | R04 | **2.63 x10<sup>-4</sup>** | 0.00574 | n.s. | 0.00871 |
|  | R10 | 0.00330 | 0.00851 | n.s. | n.s. |
|  | R15 | n.s. | 0.0433 | n.s. | n.s. |
| *Cab025935.1* | G01 | 6.85 x10<sup>-5</sup> | **8.02 x10<sup>-5</sup>** | n.s. | 0.00167 |
| *BnaA04g15260D* | G02 | **1.60 x10<sup>-4</sup>** | 4.26 x10<sup>-4</sup> | 8.46 x10<sup>-4</sup> | 3.60 x10<sup>-4</sup> |
| *BnaELF3.A04* | F01 | **1.30 x10<sup>-4</sup>** | 0.00278 | n.s. | 0.00292 |
| *AT2G25930* | F02 | n.s. | 0.0227 | n.s. | n.s. |
| Yes | R01 | 5.88 x10<sup>-4</sup> | 0.0103 | n.s. | 0.00736 |
|  | R04 | 0.00169 | 0.0129 | n.s. | 0.00611 |
|  | R10 | 0.00715 | 0.0376 | n.s. | n.s. |
|  | R15 | n.s. | 0.0459 | n.s. | n.s. |
| *Bo8g102450.1* | G01 | 1.28 x10<sup>-5</sup> | **5.45 x10<sup>-6</sup>** | **5.73 x10<sup>-6</sup>** | **2.10 x10<sup>-4</sup>** |
| *BnaC08g36440D* | G02 | 0.00930 | **2.56 x10<sup>-5</sup>** | n.s. | **3.91 x10<sup>-6</sup>** |
| *BnaAXR1.C08* | F01 | 0.00134 | **1.07 x10<sup>-4</sup>** | n.s. | **1.55 x10<sup>-5</sup>** |
| *AT1G05180* | F02 | 0.0351 | **2.79 x10<sup>-4</sup>,**<br>**#1** | 0.0201 | 0.0157 |
| No | F06 | 0.0113 | 0.0335 | n.s. | n.s. |
|  | R01 | **2.62 x10<sup>-5</sup>** | **1.67 x10<sup>-4</sup>** | n.s. | 6.20 x10<sup>-4</sup> |
|  | R04 | **2.03 x10<sup>-5</sup>** | **1.87 x10<sup>-4</sup>** | n.s. | 8.24 x10<sup>-4</sup> |
|  | R06 | 0.0133 | n.s. | n.s. | n.s. |
|  | R10 | 0.00154 | 0.00570 | n.s. | n.s. |
|  | R11 | **0.00653** | 0.00285 | n.s. | 0.0346 |
| *Cab001440.1* | G01 | 0.00144 | 0.00142 | n.s. | 0.0458 |
| *BnaA03g39820D* | G02 | 0.0170 | 0.0327 | 0.0323 | 0.00932 |
| *BnaFUL.A03* | F01 | 0.00479 | n.s. | n.s. | 0.0429 |
| *AT5G60910* | R01 | 0.00324 | n.s. | n.s. | n.s. |
| Yes | R04 | 0.007341 | n.s. | n.s. | n.s. |
|  | R06 | **0.00437, #3** | 0.0176 | n.s. | n.s. |
|  | R09 | 0.0149 | n.s. | n.s. | n.s. |
|  | R10 | **1.80 x10<sup>-5</sup>, #4** | 0.00134 | n.s. | n.s. |
|  | R15 | n.s. | 0.0088 | n.s. | n.s. |
| *Bo5g078770.1* | G01 | n.s. | 0.0183 | n.s. | n.s. |
| *BnaCnng45490D* | R02 | **0.0210, #1** | n.s. | n.s. | n.s. |
| *BnaVRN2.C05* | R03 | 0.0153 | n.s. | n.s. | n.s. |
| *AT4G16845* | R08 | **0.00769, #3** | n.s. | n.s. | n.s. |
| Yes | R09 | **0.00242, #1** | n.s. | n.s. | n.s. |
|  | R10 | 0.00478 | 0.0228 | n.s. | n.s. |
| *Cab019414.1* | G01 | 0.0198 | 0.00303 | n.s. | n.s. |
| *BnaAnng19140D* | G02 | 0.0103 | n.s. | n.s. | 0.0360 |
| *BnaVRN2.A04* | G01 | 0.00945 | n.s. | n.s. | n.s. |
| *AT4G16845* | R01 | 0.0253 | n.s. | n.s. | n.s. |
| Yes | F02 | 0.0351 | n.s. | n.s. | n.s. |
|  | R03 | **0.00784, #4** | n.s. | n.s. | n.s. |
|  | R04 | 0.0306 | n.s. | n.s. | n.s. |
|  | R08 | 0.0222 | n.s. | n.s. | n.s. |
|  | R09 | **0.00503, #4** | n.s. | n.s. | n.s. |
|  | R10 | 0.00451 | n.s. | n.s. | n.s. |
|  | R15 | 0.0269 | 0.0269 | n.s. | n.s. |
| *BnaA04g13710D* | G01 | 0.0112 | 0.0427 | n.s. | n.s. |
| *BnaA04g13710D* | G02 | n.s. | 0.0484 | 0.0225 | 0.0114 |
| *BnaCESA8.A04* | F01 | n.s. | n.s. | n.s. | n.s. |
| *AT4G18780* | R14 | **0.00639, #1** | n.s. | n.s. | n.s. |
| Yes |  |  |  |  |  |
| *Cab001225.1* | G01 | 0.00119 | **1.03 x10<sup>-4</sup>** | n.s. | 4.60 x10<sup>-4</sup> |
| *BnaA03g37880D* | G02 | 0.00236 | 0.00287 | 0.00262 | 3.98 x10<sup>-4</sup> |
| *BnaBRAVE.A03* | F01 | 5.45 x10<sup>-4</sup> | 0.00167 | n.s. | 4.09 x10<sup>-4</sup> |
| *AT2G04850* | F02 | n.s. | 0.0106 | n.s. | 0.0431 |
| Yes | R01 | 5.68 x10<sup>-4</sup> | 0.00936 | n.s. | 0.0141 |
|  | R02 | 0.0378 | n.s. | n.s. | n.s. |
|  | R03 | 0.0203 | n.s. | n.s. | n.s. |
|  | R04 | 0.00248 | 0.0172 | n.s. | 0.0184 |
|  | R06 | 0.00684 | n.s. | n.s. | n.s. |
|  | R09 | 0.00903 | n.s. | n.s. | n.s. |
|  | R10 | **1.80 x10<sup>-5</sup>,**<br>**#3** | **2.32 x10<sup>-4</sup>** | n.s. | n.s. |
|  | R15 | 0.0213 | **5.23 x10<sup>-4</sup>** | n.s. | n.s. |

Several genes appeared related to a large number of traits, as expected from high covariance between many traits (Table 2, Supplemental Table S6). These loci included two homologues of *FLC*, a gene well established as controlling flowering in Arabidopsis and Brassica species (*FLC.C02, BnaC02g00490D,* and *FLC.A03b, BnaA03g13630D*) (Raman et al., 2016; Whittaker and Dean, 2017; Schiessl et al., 2019; Song et al., 2020; Lu et al., 2022). *BnaC04g53290D*, an orthologue of *SOC1,* is closely linked to variation in all four floral initiation traits at different temperatures (R01, G01, G02/F01, Table 2). *SOC1* promotes the floral transition and this particular orthologue is upregulated in response to vernalization in winter OSR (Matar et al., 2021). These *a priori* candidates support the validation of the AT.

We found other repetitive candidates that may be causative. Examples include orthologues of; *AT1G80340* (*GA3ox2*), an enzyme involved gibberellin GA_3_, synthesis which promotes flowering (Rood et al., 1989); the floral repressor *TARGET OF EAT2* (TOE2, AT5G60120; Aukerman and Sakai, 2003); and *EARLY FLOWERING3* (*ELF3)*, a component of the evening complex of the Arabidopsis circadian clock that functions as a thermosensor (Zagotta et al., 1996; Jung et al., 2020). These genes are all floral-timing genes in Arabidopsis, however *BnaC08g36440D* is similarly linked to phenology traits (Table 2) but shows similarity to *AUXIN RESISTANT1* (*AXR1; AT1G05180*), a gene with roles in auxin signalling (Leyser et al., 1993). These repetitive candidates may be directly causally linked to all of these traits, may be highly represented within particular crop types, or may trigger initial variation in timing that then cascades through development.

Other *a priori* candidates affect speed and morphology traits. An orthologue of the floral-network and fruit tissue identity factor *FRUITFULL* (*FUL, BnaA03g39820D)*(Pajoro et al., 2014) is closely linked to extension and height traits (R06 and R10, Table 2). *AtVERNALIZATION2* (*VRN2*) is a core component of the highly-conserved histone methylase Polycomb Repressive Complex 2 (PRC2) required for vernalization at *FLC* in Arabidopsis (Whittaker and Dean, 2017; Xu and Chong, 2018). In *B. napus* we find two putative *VRN2* orthologues closely associated with all rates of raceme extension post-cold at 5°C (*BnaCnng45490D*, *BnaAnng19140D*), which may be linked to the functions of VRN2 in vascular patterning (Lucas et al., 2016). An orthologue of a regulator of growth rate, *CELLULOSE SYNTHASE 8 (BnaCESA8.A04, BnaA04g13710D*) is the most highly ranked candidate for the time between buds being visible to the start of raceme emergence at 5°C (Table 2).

### Presence/absence variation, haplotype analysis and eGWAS suggest causative variation

In order to identify specific DNA variants that could be causing phenotypic effects, we generated high-quality sequence information for a subset of *a priori* candidate genes by exome capture, and interrogated existing sequences produced by others (Chalhoub et al., 2014; Schiessl et al., 2017; Sun et al., 2017; Lee et al., 2020; Song et al., 2020; Steuernagel et al., 2021; Table 2). These candidates included two genes underneath GWAS peaks; *BnaAGL24.A01* (*BnaA01g13920D*, G03 at 10v5) and *BnaAGL15.A03* (*BnaA03g04490D*, R11 at 10°C). In some genotypes, the exome capture returned no sequence for *BnaAGL24.A01,* suggesting it may be absent. This potential presence/absence variation (PAV) is significantly linked to trait G03 for the 10v5 comparison (p = 0.0181, for fixed effect of presence, LMM with crop type as a random effect, Figure S8), although the absence variant only occurs in some spring OSR accessions. For *Bna.AGL15.A03* there was no obvious PAV, so instead we use a haplotype approach to classify the existing sequence variation (Supplemental Tables S10, S11); and found that different haplotypes were linked to variation in R11 at 10°C (p = 0.0266 for fixed effect of haplotype, LMM with crop type as a random effect, Figure S8).

Identification of DNA variants underlying AT results is complicated by the many pathways through gene regulatory networks. However, in Arabidopsis, much causative variation in expression of *FLC* is due to the presence of non-coding SNPs at the *FLC* locus itself (Whittaker and Dean, 2017; Hepworth et al., 2020). We therefore ran the transcriptome values for the candidate genes as expression GWAS (eGWAS), to identify genes regulated by their own loci (results in Supplemental Table S8). In most cases, any significant markers mapped away from the target gene. For *BnaFLC.C02,* although some GWAS models (GLM, BLINK) did produce markers on chromosome C02, few were close to the locus. However, as multiple haplotypes are known to confound GWAS approaches for *FLC* in Arabidopsis (Atwell et al., 2010), we interrogated the sequences of *BnaFLC.C02* and found three major haplotypes of *BnaFLC.C02* within the samples, as well as presence/absence variation (Supplemental Tables S10, S11). Different haplotypes and absence of *BnaFLC.C02* were associated with *BnaFLC.C02* expression at the time of measurement, though not directly with traits themselves (p = 2.298×10^−6^ and p > 0.2 respectively for fixed effect of haplotype, Linear Mixed Models (LMM) with crop type as a random effect; Figure S9).

Two SNPs were significant for *BnaELF3.A04* expression, and that with the lowest p value is within *BnaELF3.A04* itself. Presence of this marker was significantly related to *BnaELF3.A04* expression and weakly also to the G02 trait directly (p = 2.571×10^−15^ and p = 0.0775 for fixed effect of marker Cab025935.1:1608 on expression and on trait respectively, Figure 6B-C). Clustering *BnaELF3.A04* sequences resulted in three haplotype groupings that significantly related to differences in expression (p = 0.0103 for fixed effect of *BnaELF3.A04* haplotype on *BnaELF3.A04* expression, Figure 6D). There were only two non-synonmous SNPs in these haplotypes: although though one of these was in the polyQ region (Jung et al., 2020), they did not affect the predicted prion-like domain (Lancaster et al., 2014). Differences within the putative 3’UTR and to the sequence upstream of the start codon between the Darmor-*bzh* and Zhongshuang11 reference genomes may correspond to the observed expression differences.

**Figure 6:**
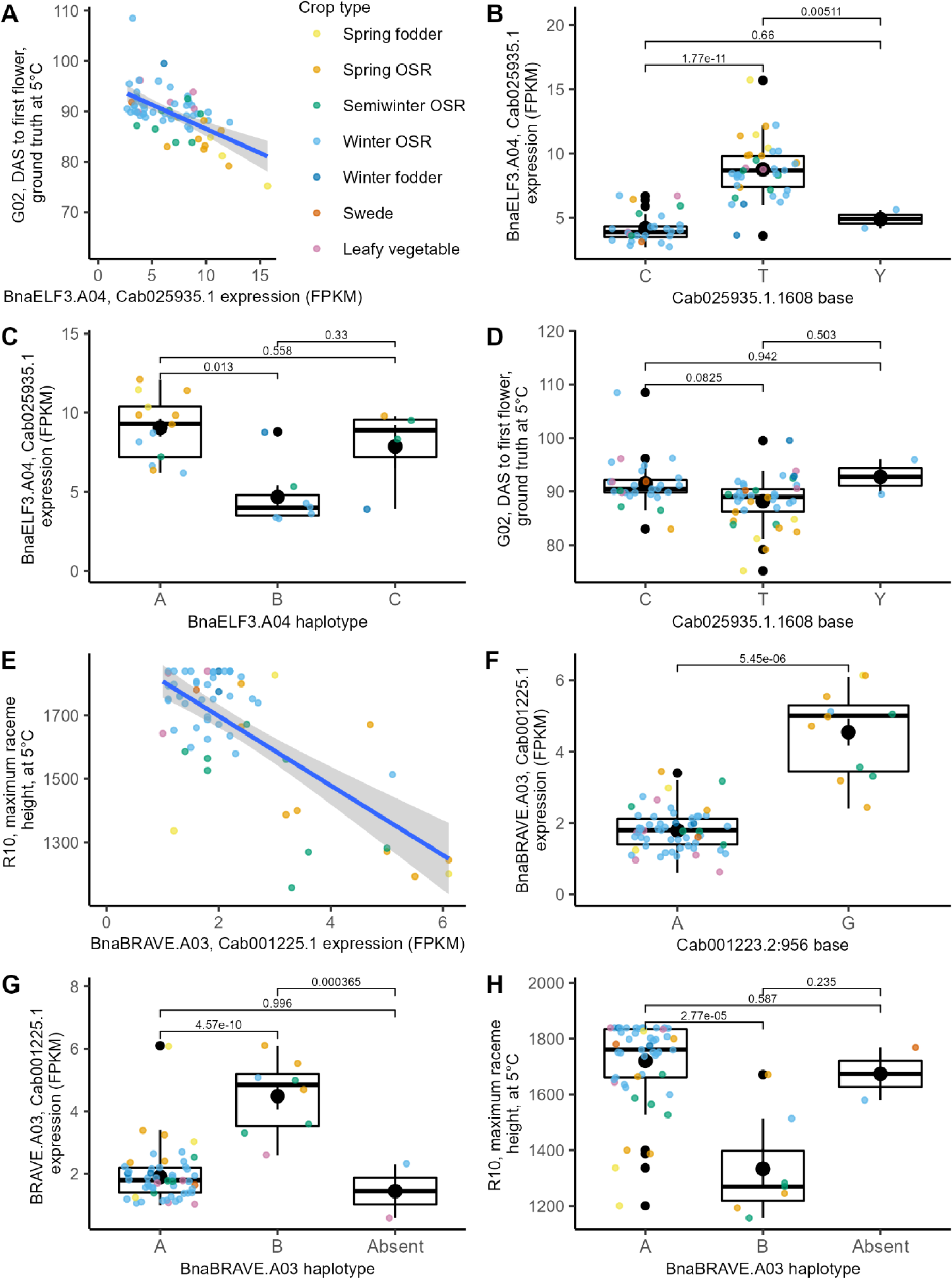
Genetic variation at BnaELF3.A04 and BnaBRAVE.A03 are associated with timing and raceme extension traits. Expression is linked to trait (A, E), and particular SNPs close to the gene are linked to its expression (B, F), and haplotypes (C, G) and traits are linked to the SNPs (D) or haplotypes (H) for *BnaELF3.A04* (A-D) and *BnaBRAVE.A03* (E-H). Individual points are estimated marginal means per accession. A, E, line is regression. B-D, F-H, boxplots showing mean (black circles) plus standard error (black lines), p values from FDR corrected post-hoc T tests.

Our exome-capture candidates also included an uncharacterised gene, *Cab001225.1*, that in a published transcriptome time-series has characteristics of downregulation similar to *FLC* during vernalization (Calderwood et al., 2021a). We included this gene to test our abilities to identify potential novel loci, and named it *BRASSICA RACEME AND VERNALIZATION* (*BnaBRAVE.A03; BRV*). In the AT, *BnaBRAVE.A03* linked to buds visible at 10°C and maximum raceme height at both temperatures, and more weakly to raceme growth rate and duration traits (Table 2, Figure 6E). Four significant GWAS peaks were associated with *BnaBRAVE.A03* eGWAS. The most significant located to chromosome C03, not close to the homeologue *BnaBRAVE.C03*, but in a region of the Darmor-*bzh* genome with synteny to the Arabidopsis *BRAVE* region, which may therefore harbour a *BRAVE* copy in some accessions. The second association is close to splicing regulator *FPA,* while the fourth is in *Cab035024.1*, or *MYB13* (*AT1G06180*), a regulator of inflorescence growth (Kirik et al., 1998). The third highest hit is in *Cab001223.2*, two gene models away from *BnaBRAVE.A03* (p = <2×10^−16^ for fixed effect of marker on *BnaBRAVE.A03* expression, Figure 6F). *BnaBRAVE.A03* has both haplotype structure and presence/absence variation (PAV), which are significantly linked both to expression of the gene and the linked phenotype (p = 3.065×10^−9^ and p = 8.40×10^−5^ for fixed effect of *BnaBRAVE.A03* on *BnaBRAVE.A03* expression and on R10 maximum height after 10°C respectively, Figure 6G, H).

### A mutant in *B. rapa BRAVE* shows altered floral raceme outgrowth

To further assess whether *BRAVE* may be involved in raceme extension, we obtained a mutant in *BRAVE.A03* in a *Brassica rapa* oilseed, R-o-18 (Stephenson et al., 2010). *Brassica rapa* is the ancestral parent species of the *B. napus* A genome, so *BraBRAVE.A03* and *BnaBRAVE.A03* are likely to share function. *BraBRAVE.A03* is 403 residue protein predicted to contain dopamine beta-monooxygenase (DOMON) and membrane-integral cytochrome b domains. The mutant line has a G>A conversion creating a stop codon at residue 226, disrupting the cytochrome b561 domain (Supplemental Table S11). We genotyped a segregating F2 population from a second backcross to the unmutagenised parent. R-o-18 is a rapid-cycling ‘spring’ type, and when grown without vernalization plants containing the *Brabrave.a03* allele were slightly, but not significantly slower to BBCH51 or first flower, and showed no change in main raceme height (Student’s T p > 0.2, Figure 7A-B). However, in *B. rapa*, *Brabrave.a03* racemes converted to flower production later in development, producing two more leaves before the floral transition than the wild type (Student’s T p = 0.00969; 16.7 leaves mean; 14.9 leaves mean, Figure 7C). We compared this result with a manual count of leaves produced in the *B. napus* experiment, and found that after both temperature treatments, absence of *BnaBRAVE.A03* was also associated with a significant developmental delay in the floral transition (for 10°C, p = 0.0155, for 5°C, p = 0.0418, post-hoc T-tests on LMM, Figure 7E-F).

**Figure 7:**
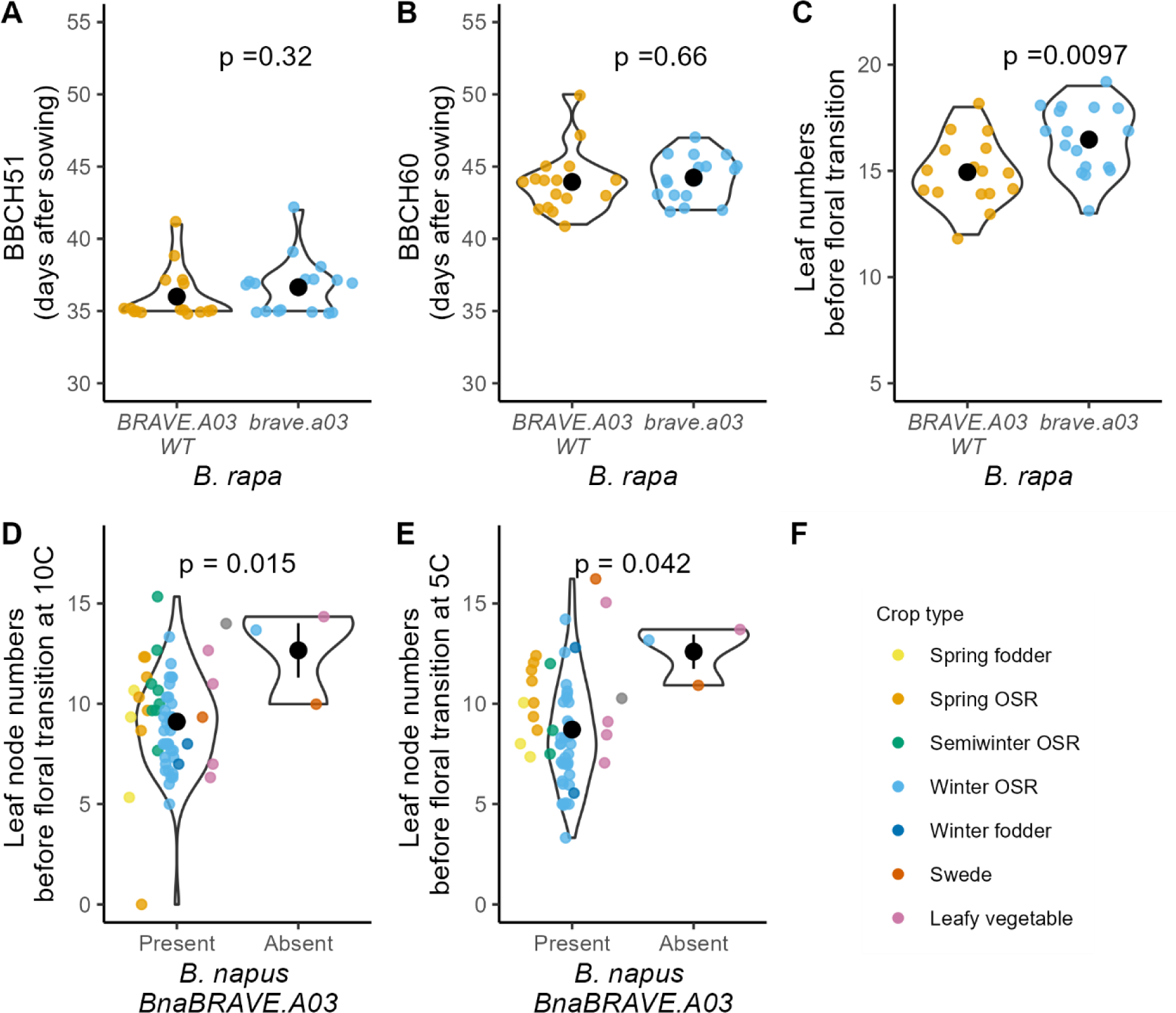
BRAVE.A03 is linked to the floral transition in *B. rapa* and *B. napus*. Measurement of calendar (A, B) and developmental (C-E) time to inflorescence production by presence of a full copy of *BRAVE.A03*. A-C) Wild-type and mutant homozygotes from *B. rapa* backcrossed F2 population phenotyped for floral buds visible (BBCH51; A); first flower opening (BBCH60; B); and number of leaves on the main raceme before floral transition. N = 17 *Brabrave.a03*; n = 16 *BraBRAVE.A03*. P values are Student’s T. D-E) Number of leaves before floral transition, manually counted, for *B. napus* plants from phenomics experiment, individual points are estimated marginal means per accession, coloured by crop type. P-values are post-hoc Student’s T pairwise contrasts (see Methods). Violin plots showing mean (black circles) plus standard error (black lines).

## Discussion

### Study of inflorescence responses to vernalization temperature could contribute to breeding for warmer winters

Vernalization response is a key delimiter of crop type within the diverse *Brassica napus* species. The effect of vernalization on bolting and flower initiation has been well characterised using traditional manual methods and these studies have revealed much of the genetic and molecular architecture underlying the observed phenotypic variation (for example O’Neill et al., 2019; Schiessl et al., 2019; Song et al., 2020; Calderwood et al., 2021a). Unlike the visually distinctive developmental phase change that results in the opening of the first flower, subsequent phenological stages, such as flower duration and number, are much more difficult to quantify using manual approaches. Thus, few population scale studies have attempted to measure variation in these traits. This generates an important gap in our knowledge given their importance to yield (Kirkegaard et al., 2018; Siles et al., 2021). The yield of winter OSR in the UK is reduced by warmth in early winter (Brown et al., 2019; O’Neill et al., 2019) and this can only partly be explained by a delayed floral transition (Lu et al., 2022). In this study, recurrent imaging of multiple *B. napus* plants captured a detailed longitudinal record of each plant’s development, permitting the isolation of multiple digital phenotypes characterising flowering and raceme development. In line with previous observations, we find that warmer vernalization leads to a late floral transition (O’Neill et al., 2019), However, our work also demonstrates that vernalization at higher temperature leads to slower growing and shorter racemes with lower flowering area for winter OSR, perhaps contributing to the observed reduction in yield. This is also largely true for spring OSR accessions, which would be presumed to be impeded by lower growing temperatures. However, for semiwinter OSR, we observed a higher pre-peak total flowering area under the warmer vernalization conditions, providing a rationale for its yield in warmer climates.

### High throughput imaging potentiates accurate measurement of novel traits

Our automated image capture measurements were highly repeatable between replicates, and manual data agreed very well with the digital equivalent. These two factors provided us with the confidence that our trait extraction procedures were robust and capable of quantifying variation across time, between genotypes and in response to the treatments.

Most previous Brassica high-throughput imaging studies have been of smaller scale, lower dynamic resolution, or undertaken earlier in development and limited to modelling biomass accumulation and height measurements (e.g. Knoch et al., 2020; Li et al., 2020). Only recently have methods started to appear with relevance to quantification of flowering (Ebersbach et al., 2022), and post transition phenotypes (Ghahremani et al., 2021), but their population size has been too restricted to allow GWAS. Platform capacity remains a key limitation for this technology. Another limitation is the frequency of imaging. Locally this is restricted to once per day when the platform is full. With more frequent image collection, e.g. two to four times a day, it is possible to measure diurnal growth features (Faralli et al., 2019), though sacrificing population size. Moreover, image resolution may be limiting for accurately measuring smaller features, such as pods. Higher resolution images using bespoke camera configurations have the potential to provide significantly more information but at the cost of higher computational and data storage requirements, particularly if combined with 3D modelling and multispectral imaging.

Nevertheless, the images acquired in this experiment have potential for further analysis. Here we have used daily metadata to calculate first order raceme extension. Further phenotypes could also be modelled by analysing time-stamped images and watering events with respect to environmental data and periods of abiotic stress, such as low relative humidity. Additional post-floral transition features likely to impact on crop yield include stem diameter, branching angle, flower shape & persistence as well as fertilisation and aspects of pod development (Hamidinekoo et al., 2020; Ghahremani et al., 2021). These traits are outside the scope of the current study, but we have made the BR017 image dataset available (doi: <u>10.20391/35c7b57c-09a4-43e5-a7d7-89fb4f420be0</u>) to encourage data re-use and the development of additional protocols, such as AI deep learning (Hamidinekoo et al., 2020).

Even without further analysis, processing the large number of resulting association models became a bottleneck. As this is likely to be a recurrent problem as High Throughput Phenotyping studies produce increasingly more data through computational methods (Li et al., 2020; Ghahremani et al., 2021; Ebersbach et al., 2022), we developed and exploited an automated pipeline to run both GWAS and AT for all the generated traits (Nichols et al., in prep.), saving research time.

Clear genetic signatures for *a priori* candidates, such as *FRUITFULL* and *CELLULOSE SYNTHASE 8* for raceme characteristics, support the biological relevance of the novel dynamic traits. High time resolution increases power to detect loci that make small gains (Camargo et al., 2016; Knoch et al., 2020), and to further dissect the complexity of interrelated ‘final’ phenotypes through pathway analyses (Calderwood et al., 2021b). Identifying these time-critical effects on development holds promise for the search for unexploited natural variation to break trade-offs and improve crop productivity.

### Measurement of dynamic growth reveals candidate regulators throughout development

The rich trait information generated was used to undertake GWAS and AT to identify candidate genes contributing to crop type differences. Although the population size tested here was very small (69 or fewer accessions were used for GWAS, depending on the trait), it was sufficient to identify highly significant GWAS peaks for several traits, including those over promising variants of *AGAMOUS-LIKE* genes. Further, via AT approaches we were able to identify potentially causative variation in *a priori* genes. For example, multiple orthologues of *FLC* are major contributors to crop type differences in flowering time control (Schiessl et al., 2019; Wu et al., 2019; Song et al., 2020; Calderwood et al., 2021a). One of the primary candidates revealed by this study is *BnaFLC.C02*, previously identified as mainly important for variation in spring material when vernalised at 5°C (Schiessl et al., 2019; Schiessl, 2020). Here the relative contribution of this gene was highest in the warmer temperature treatment, a conclusion supported in field-like experiments (Lu et al., 2022), confirming the efficacy of our pipeline and suggesting that deletion variants might be valuable targets for breeding for warmer winters.

Potentially causative DNA variation may also be identified via AT by using eGWAS, to find variation that is *cis-*encoded at both *a priori* and previously unidentified candidates, as we show here for *BnaELF3.A04* and *BnaBRAVE.A03*. The expression and protein activity of *ELF3* are known to influence flowering, so the haplotype structures identified here might have effects on both aspects of regulation (Zagotta et al., 1996; Jung et al., 2020). As an uncharacterised gene, the effects of the differences between the *BnaBRAVE.A03* haplotypes are difficult to predict, but incomplete coding variants lead to a delayed floral transition in both *B. napus* and *B. rapa*, suggesting that *BRAVE.A03* is linked to promotion of inflorescence development. A related gene in *Brassica carinata*, *COPPER INDUCED in LEAVES1 (BcCIL1*) is involved in redox control of lateral root growth downstream of auxin and abscisic acid signalling (Gibson et al., 2012), while in *Medicago falcata* the *CIL1* orthologue, *AUXIN INDUCED IN ROOT CULTURES12 (AIR12),* is reported to be involved in cold tolerance (Wang et al., 2021). *Bna-* and *BraBRAVE.A03* lack the GPI-anchor domain of these proteins. Its closest Arabidopsis orthologue (*At2g04850*) is an outlier to the rest of the *AIR12* family (Preger et al., 2009), but has previously been suggested as a candidate for a QTL for root growth responses to phosphate in *B. napus* (Zhang et al., 2016). Our analysis highlights this gene for further mechanistic investigation.

In summary, we present here a longitudinal approach to study development with high throughput phenotyping. Even in a relatively small population, we demonstrate how this approach allows us to identify known regulators of well-characterised traits as well as unknown regulators and previously unmeasurable traits, potentiating variant and gene discovery.

## List of author contributions

KW, FC, LØ, RJM, JHD, RW designed the research; KW, BS, FC, RW performed research; KW, BN, HW, PP, BS, contributed new computational tools; KW, JH, BN analysed data; KW, JH, BN, FC, JHD, RW wrote the paper. No natural language processing tools (e.g. ChatGPT) were used during the course of this work.

## Acknowledgments

The authors would like to acknowledge the huge contribution to this work of Judith Irwin, now retired. We would like to acknowledge useful discussions with Lenka Havlickova, Zhesi He and Ian Bancroft in preparation for the GWAS, AT and targeted sequence capture work, and Alexander Calderwood for gene expression data exploration. Many thanks to Steve Penfield for comments that improved the manuscript substantially. Thanks to Saleha Bahkt, Daniella Shipley, Catherine Taylor, Richard Goram and Sophie Preston of the JIC technology platforms for plant materials, care and genotyping. We would also like to acknowledge the work of many colleagues on GWAS and flowering in *B. napus* that we did not have space to cite.

## Materials and Methods

### Plant materials and growth conditions

The experiment was run across two compartments of an environmentally controlled glasshouse equipped with a semi-automated conveyor system (LemnaTec Scanalyzer). Images were acquired using the modified RGB cabin of the platform. Full description of the parameters used for the experiments is in Supplementary Methods.

Longitudinal image data were collected from 72 genotypes exposed to two different vernalization treatments. 71 genotypes were from the RIPR panel, plus an in-house variant of the winter OSR, Darmor (ET) (Tudor et al., 2020). Effective crop type designations were adapted from data from the ASSYST wider diversity panel (Bus et al., 2011), listed in Supplemental Table S9. Seed were sown in 8 cm square pots of Levington F2 compost. Three replicates of each genotype-treatment combination were vernalised for 8 weeks at 5 °C (treatment 1) or 10 °C (treatment 2) giving a total of 426 plants. Vernalization conditions were 8 hour photoperiod with fluorescent lighting at 60 µmmol.m^−2^.s^−1^.

Post-vernalization, plants were potted on into 3.5 L square pots of Levington M3 compost. A uniform weight of compost was used in all pots to facilitate target weight watering to 70% soil water content. All plants were watered daily at the top of the pot until their roots were established (1 week), and then into the pot saucers. After four weeks, watering was switched to twice-daily to minimise plant-wilting during a period of high temperatures. Plants were paired together on the platform in an attempt to minimise local environment effects, such that replicate *n* for each genotype had treatments 1 and 2 side-by-side. The plants were pseudo-randomised by genotype, with Compartment 5 containing all replicate 1 plants plus replicate 2 plants up to genotype 36; Compartment 6 held all replicate 3 plants plus replicate 2 plants from genotype 37 to genotype 71. An extra 86 compost-only pots were interspaced to facilitate local corrections to water use calculations. Compartment settings were: 18°C day/15°C night temperature, 14 hour photoperiod, with 600 W Son-T lights to supplement daylight to 350 W.m^−2^.s^−1^. In an attempt to minimise tangling as the plants grew, an empty car was set between consecutive plants. Specimens were removed when too large for the system, and grown to full maturity off-platform. 13888 plant × day image sets were obtained between 6 June & 28 August 2018.

### BRAVE mutants

Line ji30491-a from the JIC Molecular Genetics *B. rapa* Targetting Induced Local Lesions IN Genomes (TILLING) population (www.jic.ac.uk/research-impact/technology-research-platforms/molecular-genetics; Stephenson et al., 2010) was identified as containing a stop mutation. An M3 mutant was backcrossed twice into the non-mutagenised parent line R-o-18. The mutation was genotyped using KASP markers (Supplemental Table S12). In the F2 generation, seedlings were sown in P24 trays for three weeks, genotyped, then transferred to 1 litre pots, resulting in 18 wild-type plants and 17 plants homozygous for the mutation. Plants were grown in ‘cereal mix’ (65% peat, 25% loam, 10% grit, 3 kg m^−3^ dolomitic limestone, 1.3 kg m^−3^ PG mix, 3 kg m^−3^ Osmocote Mini 16-8-11 2 mg + Te 0.02% B, 300 g m–3 Exemptor) in a glasshouse set to 16h light + 18 °C/8h dark + 15°C, with 600W lamps to provide supplementary lighting, 70% humidity.

### Computational methods

We developed processes to extract features and quantitative data as described in Table 1, Figures 1 and 2 and Supplemental Figure 3. These methods are expanded in Supplementary Methods.

Four steps are used in the pipeline to derive the flowering features listed in Table 1;

- segmentation of native side view images
- checks for erroneous segmentation
- digital phenotype calculations for plants
- digital phenotype statistics for populations

Overhead images were used in a similar fashion to the side view images for evaluating flowering onset only.

An initial segmentation routine was developed to build a simple binary mask of each RGB plant image, with all background noise removed to calculate the height of each plant. Height versus time and its first order growth derivative were parametised to give the digital phenotypes R01 to R12, described in Table 1. The onset of raceme extension was defined as the interpolated timestamp 48 hours prior to the raceme passing through the 500 pixel height barrier. This definition was determined by observation. Model curves and parameter derivations are given in Supplemental Figure 3 and Supplemental Methods Table 2.

For a small number of genotypes, raceme extension and flowering had already commenced before the plants were loaded onto the LemnaTec platform; this phenotype was extracted and subsequently used as R15. Data affected by this or by raceme collapse (R13) was removed.

### Temperature responsiveness

For the 10-5 comparison, values for each trait measured on a plant treated with 5°C vernalization were subtracted from the values from its paired 10°C treated plant (see ‘Plant materials and growth conditions’). For 10v5, the same measurements were divided, except for the absolute timing traits, where the time before the end of vernalization was subtracted before division, to compare only post-treatment effects. Binary traits (R13, R15) were excluded from this analysis.

### Statistical and genome wide association analysis

Trait data in Figures 3, 4, 6, S4, S5, S6, S8 and S9 are estimate marginal means per genotype from LMMs run for each treatment separately, defining location (blocking factor) as a random effect and genotype as a fixed effect. These estimate means were analysed and run through a GEM association and a GWAS using a joint pipeline called GAGA, available at https://github.com/bsnichols/GAGA (Nichols et al., in prep.), using the Pantranscriptome database for the RIPR panel as described by Havlickova et al. (2018), version 11, available at http://www.yorknowledgebase.info/. The GAGA pipeline was conducted in R using the statistical analysis, GEM and GWAS functions. The estimate means test used in this study was the Linear Mixed Model (LMM). The GAGA pipeline uses GAPIT for GWAS (Wang and Zhang, 2021), and the three models used in this study were; GLM (General Linear Models), FarmCPU and Blink. GAGA extracts data on significant markers from all three models (Table S5) but compares QQ plots to select the best fitting model for final analysis. The plotting step produces Manhattan plots for both the GEM association and GWAS results.

25 traits were run for each temperature and temperature comparison (G01-G03, R01-R15, F01-3, F06, F07, F09-F10), with the exception of R13 and R15 for the 10°C vs 5°C analysis because these binary traits generated division by zero errors. For eGWAS, the FPKM values for the selected GEMs (Table 2) were extracted from the AT database and run as normal GWAS traits.

Linkage disequilibrium regions were calculated using TASSEL version 5.2.70 (Bradbury et al., 2007).

All other statistics were computed using R (R Core Team, 2018). P-values in Figures 3, 4, S4, S5 and S6 are pairwise contrasts from LMMs with the form: trait ~(1|location)*crop type*temperature; for Figures 6, S8, S9, are FDR corrected post-hoc test from LMMs as y ~(1|crop type)+x.

### Targeted Sequence Capture

Genomic DNA was extracted from a panel of 96 *B. napus* plants as described by Woodhouse et al. (2021). We used a bait library targeting a priori flowering time genes for sequence enrichment. The bait library is available at https://doi.org/10.5281/zenodo.4473283 (Steuernagel et al., 2021). Bait synthesis and target enrichment was performed by Arbor Biosciences (https://arborbiosci.com/). Sequencing and alignment was carried out as previously (Woodhouse et al., 2021) to the Darmor-*bzh* v4.1, Zhongshuang 11, Express 617, Westar 10 DH and Tapidor v3 reference genomes (Chalhoub et al., 2014; Sun et al., 2017; Lee et al., 2020; Song et al., 2020).

Sequences were viewed on Integrative Genomics Viewer (Robinson et al., 2011) to check for quality, presence/absence variation, and haplotype scoring. Consensus sequences were aligned with Clustal Omega and PhyML maximum likelihood trees were generated in UGENE to check haplotype scoring (Okonechnikov et al., 2012). Prion-like domains for ELF3 variants were assessed by PLAAC (Lancaster et al., 2014).

### Data

Image data is deposited at the Aberystwyth University Datasets, doi: <u>10.20391/35c7b57c-09a4-43e5-a7d7-89fb4f420be0</u>.

Raw data for targeted sequence capture experiments is available at EBI, study number PRJEB57649, https://www.ebi.ac.uk/ena/browser/view/PRJEB57649.The bait library is available at https://doi.org/10.5281/zenodo.4473283 (Steuernagel et al., 2021).

The functional genotypes are available from the York Oilseed Rape Knowledgebase (http://www.yorknowledgebase.info/).

