## Supplemental data for "Integrated Phenomics and Genomics reveals genetic loci associated with inflorescence growth in *Brassica napus*"

### Supplemental Figures

#### Supplemental Figure S1. Example of T1 and T2 flowering curves.

Red dots represent the original yellow pixel count (flower + leaf); black circles are a minimum value of {∑pixel_>50_, 5×∑pixel_>75_} and green dots are interpolated results between black circles. Each panel represents an individual replicate plant.

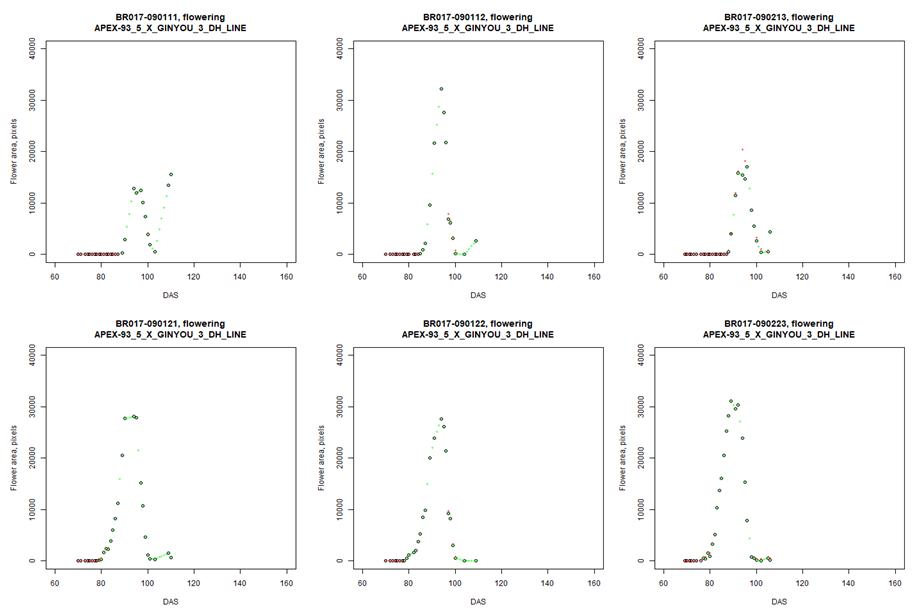

#### Supplemental Figure S2. Onset of flowering - correlation between side view image analysis and top view image analysis.

Onset of flowering, extracted from images from the side view versus top view. Outliers in the top view data were found to be due to the flowering area of the raceme bending out of the selected region of interest.

First flowers, top view, DAS

| First flowers, side view, DAS | first_flowers_DAS |
| --- | --- |
| 91 | 91 |
| 91 | 91 |
| 95 | 94 |
| 94 | 94 |
| 89 | 89 |
| 92 | 92 |
| 90 | 90 |
| 92 | 92 |
| 97 | 95 |
| 96 | 96 |
| 89 | 89 |
| 92 | 92 |
| 89 | 89 |
| 89 | 89 |
| 90 | 90 |
| 89 | 89 |
| 86 | 87 |
| 85 | 86 |
| 89 | 89 |
| 85 | 85 |
| 79 | 79 |
| 79 | 79 |
| 88 | 89 |
| 77 | 77 |
| 90 | 90 |
| 94 | 104 |
| 94 | 94 |
| 91 | 91 |
| 89 | 89 |
| 89 | 89 |
| 94 | 94 |
| 90 | 90 |
| 94 | 94 |
| 94 | 94 |
| 88 | 88 |
| 91 | 91 |
| 89 | 89 |
| 89 | 89 |
| 90 | 90 |
| 89 | 89 |
| 87 | 87 |
| 87 | 87 |
| 94 | 94 |
| 91 | 91 |
| 94 | 94 |
| 94 | 94 |
| 91 | 91 |
| 91 | 91 |
| 94 | 94 |
| 94 | 94 |
| 97 | 97 |
| 95 | 95 |
| 105 | 105 |
| 92 | 92 |
| 91 | 91 |
| 90 | 90 |
| 91 | 91 |
| 92 | 92 |
| 89 | 88 |
| 89 | 89 |
| 91 | 91 |
| 91 | 91 |
| 94 | 94 |
| 91 | 91 |
| 89 | 88 |
| 89 | 89 |
| 94 | 94 |
| 91 | 91 |
| 91 | 91 |
| 89 | 91 |
| 91 | 95 |
| 88 | 88 |
| 94 | 94 |
| 94 | 94 |
| 102 | 102 |
| 99 | 100 |
| 91 | 91 |
| 99 | 99 |
| 97 | 97 |
| 99 | 98 |
| 118 | 123 |
| 117 | 117 |
| 94 | 94 |
| 119 | 119 |
| 100 | 100 |
| 94 | 94 |
| 102 | 102 |
| 100 | 100 |
| 95 | 95 |
| 98 | 98 |
| 97 | 97 |
| 91 | 91 |
| 94 | 94 |
| 89 | 89 |
| 88 | 88 |
| 91 | 91 |
| 94 | 94 |
| 92 | 92 |
| 95 | 95 |
| 94 | 94 |
| 89 | 89 |
| 94 | 94 |
| 89 | 89 |
| 89 | 89 |
| 91 | 91 |
| 89 | 89 |
| 87 | 87 |
| 88 | 88 |
| 94 | 94 |
| 92 | 91 |
| 101 | 100 |
| 100 | 100 |
| 91 | 91 |
| 105 | 105 |
| 94 | 94 |
| 94 | 94 |
| 97 | 97 |
| 104 | 104 |
| 92 | 92 |
| 96 | 96 |
| 91 | 91 |
| 91 | 92 |
| 97 | 97 |
| 95 | 94 |
| 88 | 88 |
| 95 | 94 |
| 89 | 89 |
| 89 | 89 |
| 94 | 94 |
| 94 | 92 |
| 87 | 87 |
| 91 | 91 |
| 90 | 90 |
| 89 | 91 |
| 94 | 94 |
| 94 | 94 |
| 91 | 91 |
| 91 | 91 |
| 91 | 91 |
| 94 | 94 |
| 97 | 97 |
| 96 | 96 |
| 91 | 91 |
| 94 | 94 |
| 94 | 94 |
| 94 | 94 |
| 97 | 97 |
| 98 | 97 |
| 92 | 92 |
| 92 | 92 |
| 95 | 94 |
| 94 | 94 |
| 109 | 102 |
| 104 | 104 |
| 98 | 98 |
| 99 | 99 |
| 91 | 91 |
| 90 | 90 |
| 94 | 94 |
| 94 | 94 |
| 89 | 89 |
| 96 | 96 |
| 95 | 97 |
| 95 | 92 |
| 90 | 90 |
| 89 | 87 |
| 95 | 95 |
| 87 | 87 |
| 101 | 101 |
| 117 | 117 |
| 130 | 130 |
| 131 | 130 |
| 125 | 137 |
| 123 | 123 |
| 91 | 91 |
| 91 | 91 |
| 102 | 102 |
| 94 | 94 |
| 88 | 88 |
| 106 | 105 |
| 89 | 89 |
| 89 | 89 |
| 90 | 90 |
| 94 | 94 |
| 89 | 89 |
| 87 | 87 |
| 75 | 75 |
| 75 | 74 |
| 73 | 78 |
| 70 | 70 |
| 74 | 73 |
| 69 | 69 |
| 82 | 82 |
| 82 | 82 |
| 71 | 70 |
| 70 | 70 |
| 86 | 89 |
| 69 | 69 |
| 86 | 89 |
| 85 | 87 |

#### Supplemental Figure S3. Computational identification of raceme characteristics.

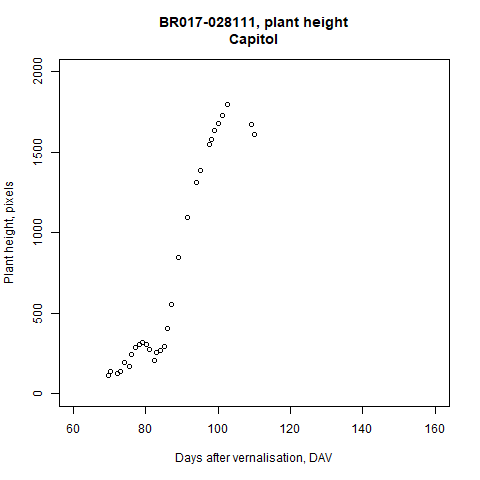
A, a sample dataset growth curve. B, a sample first order derivative curve. C, An idealised raceme growth curve showing derivation of traits R01, R05-R07, R10-R12. D, An idealised first order derivative (growth rate) curves calculated using the growth over the preceding period to give a normalised daily rate, for extraction of R02-R04, R08-R09.

A

B

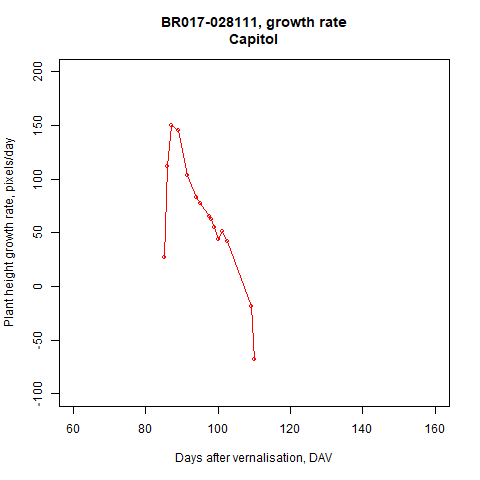

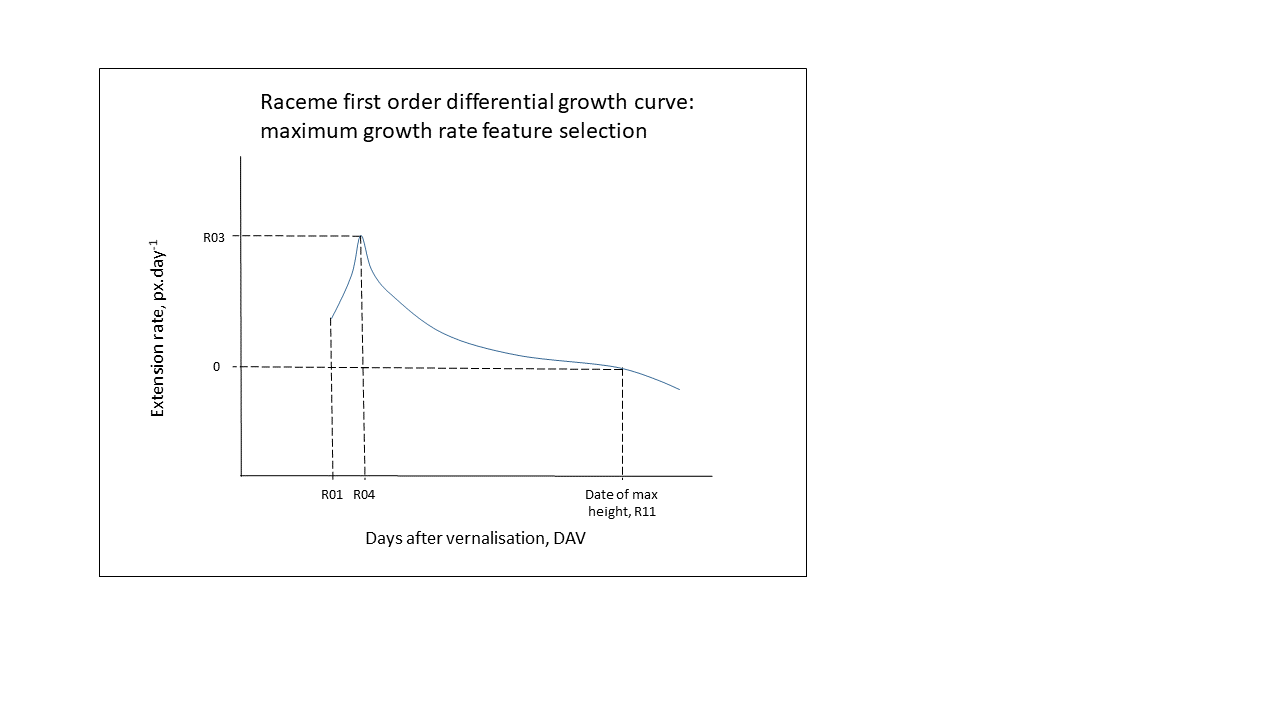

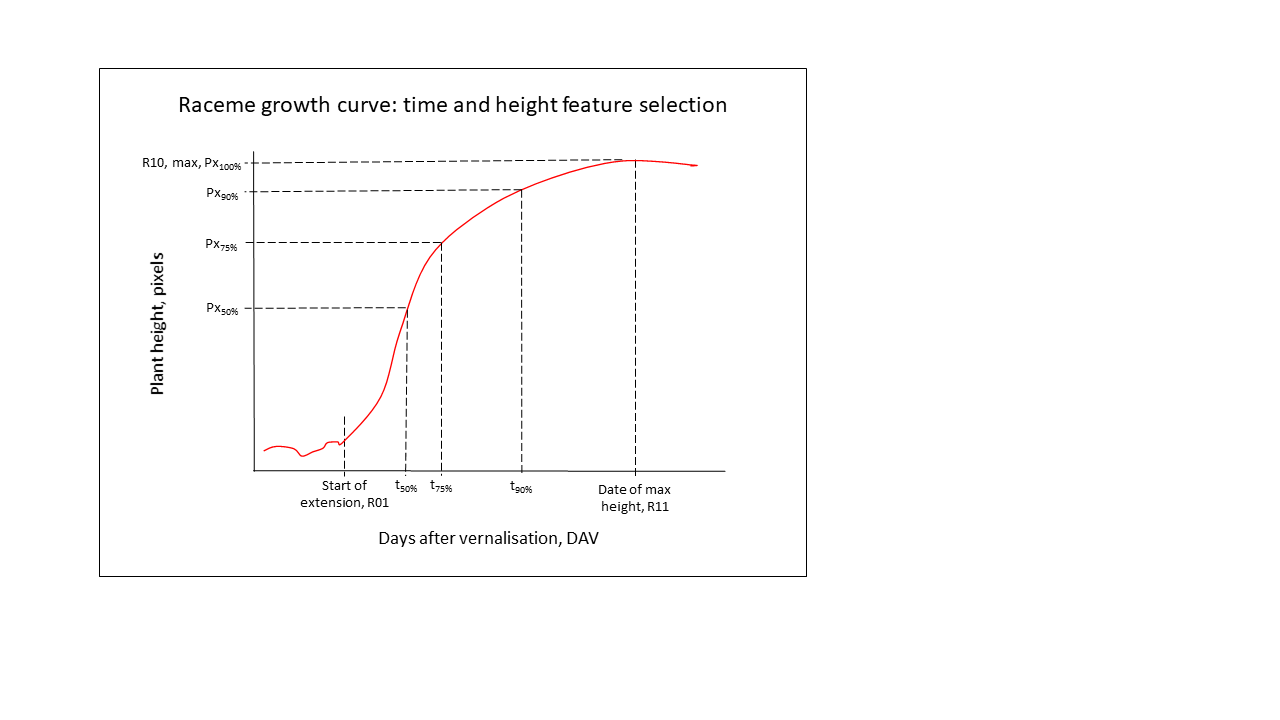

C
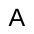

D

#### Supplemental Figure S4. Comparison of phenology responses to different vernalization temperatures.

Comparison of responses to different vernalization temperatures for: A-B) R01, days from sowing to raceme extension starting; C-D) G02, days from sowing to first flower opening by manual observation; E-F) F01, days from sowing to first flower opening by automated analysis; G-H) G03, days from manual buds visible to manual first flower scoring. Panels A, C, E, G show boxplots by crop type and vernalization treatment, p-values are Tukey-corrected pairwise contrasts per crop type for response to vernalization temperature (see Methods). Panels B, D, F, H show scatter plots with trendline showing linear regression; B) R^2^=0.779, p-value= < 2.2x10^-16^; D) R^2^=0. 759, p-value= < 2.2x10^-16^; F) R^2^=0.766, p-value= < 2.2x10^-16^; H) R^2^=0.1891, p-value=2.18x10^-4^. Points on all graphs show genotype estimated marginal means from Linear Mixed Model. N varies per treatment and trait, but maxima N are: Spring fodder = 3, spring OSR = 8, semiwinter OSR = 8, winter OSR = 42, winter fodder = 2, swede = 2, leafy vegetable = 6.

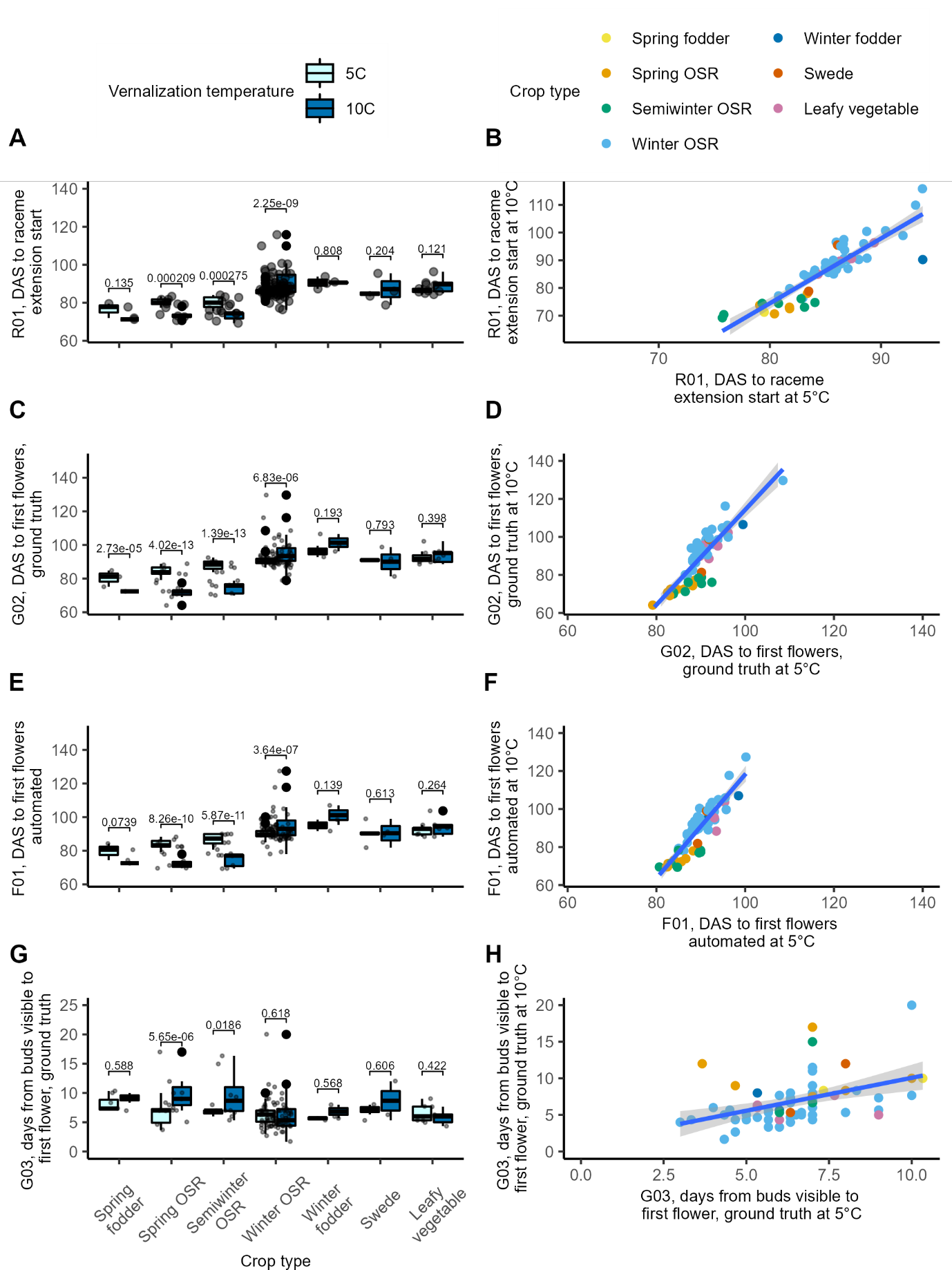

#### Supplemental Figure S5. Comparison of phenology responses to different vernalization temperatures.

Comparison of responses to different vernalization temperatures for: A-B) R14, days from buds visible (manual) to raceme extension starting (automated); C-D) R04, days from sowing to first flower opening by manual observation; E-F) F02, days after sowing to peak flowers; G-H) R11, days from sowing to first flower opening by automated analysis. Panels A, C, E, G show boxplots by crop type and vernalization treatment, p-values are Tukey-corrected pairwise contrasts for crop type response to vernalization temperature (see Methods). Panels B, D, F, H show scatter plots with trendline showing linear regression; B) R^2^=0.0953, p-value= 0.00752; D) R^2^=0. 760, p-value < 2.2x10^-16^; F) R^2^=0.640, p-value= 2.214^-15^; H) R^2^=0.708, p-value < 2.2x10^-16^. Points on all graphs show genotype estimated marginal means. N varies per treatment and trait, but maxima are: Spring fodder = 3, spring OSR = 8, semiwinter OSR = 8, winter OSR = 42, winter fodder = 2, swede = 2, leafy vegetable = 6.

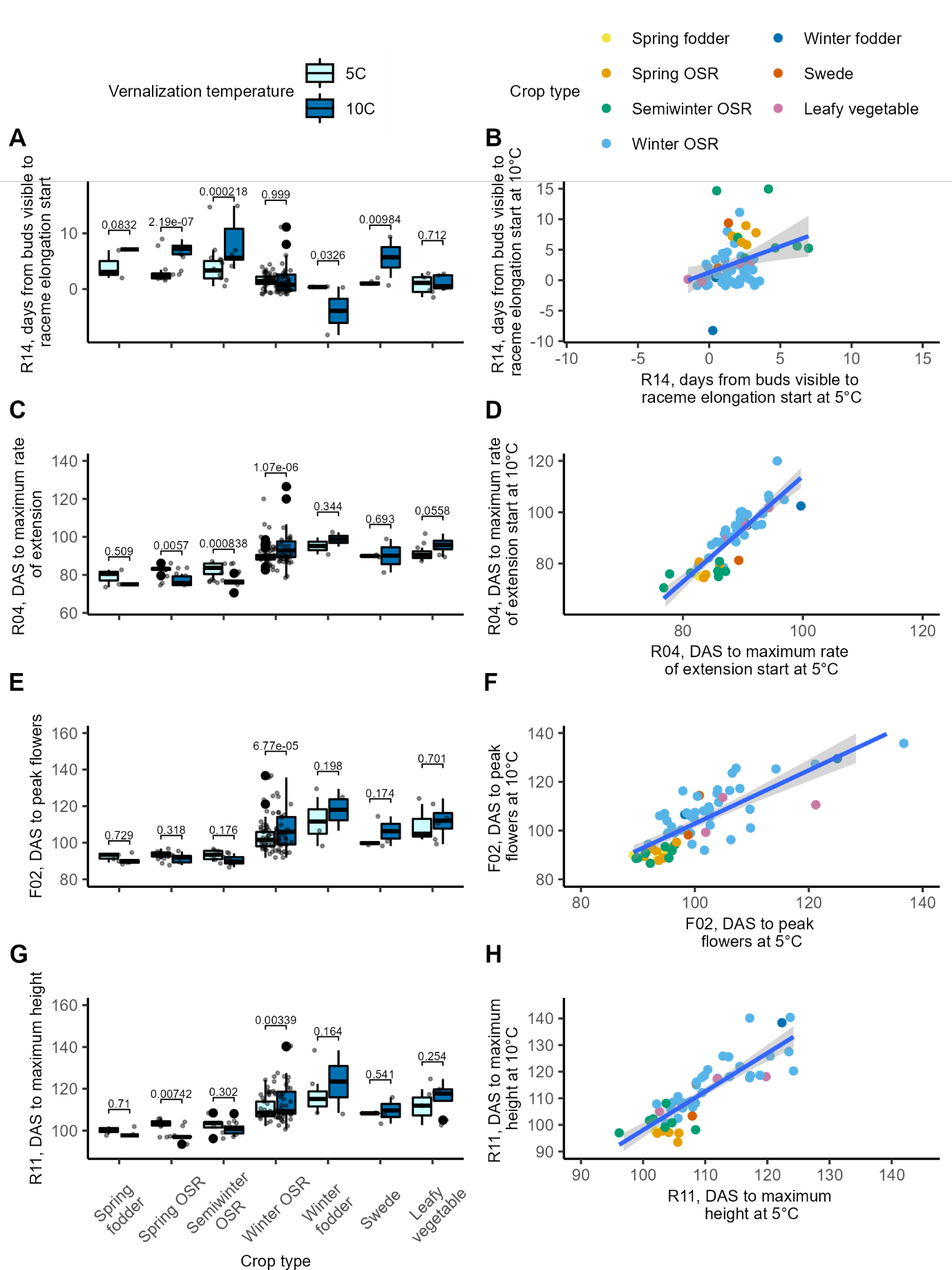

#### Supplemental Figure S6. Comparison of morphological responses to different vernalization temperatures

A) R03, maximum rate of raceme extension, in pixels per day; B) R05, main period of extension, days from the start of raceme extension to reaching 90% of the maximum height; C) R06, days from the start of raceme extension to reaching 50% of the maximum height; D) R08, mean rate of extension from the start to reaching 50% of the maximum height; E) R09, mean rate of extension from the start to reaching 75% of the maximum height; F) R12, days from raceme extension starting to maximum raceme height. Boxplots by crop type and vernalization treatment, p-values are Tukey-corrected pairwise contrasts (see Methods), points represent genotype estimated marginal means. N varies per treatment and trait, but maxima are: Spring fodder = 3, spring OSR = 8, semiwinter OSR = 8, winter OSR = 42, winter fodder = 2, swede = 2, leafy vegetable = 6.

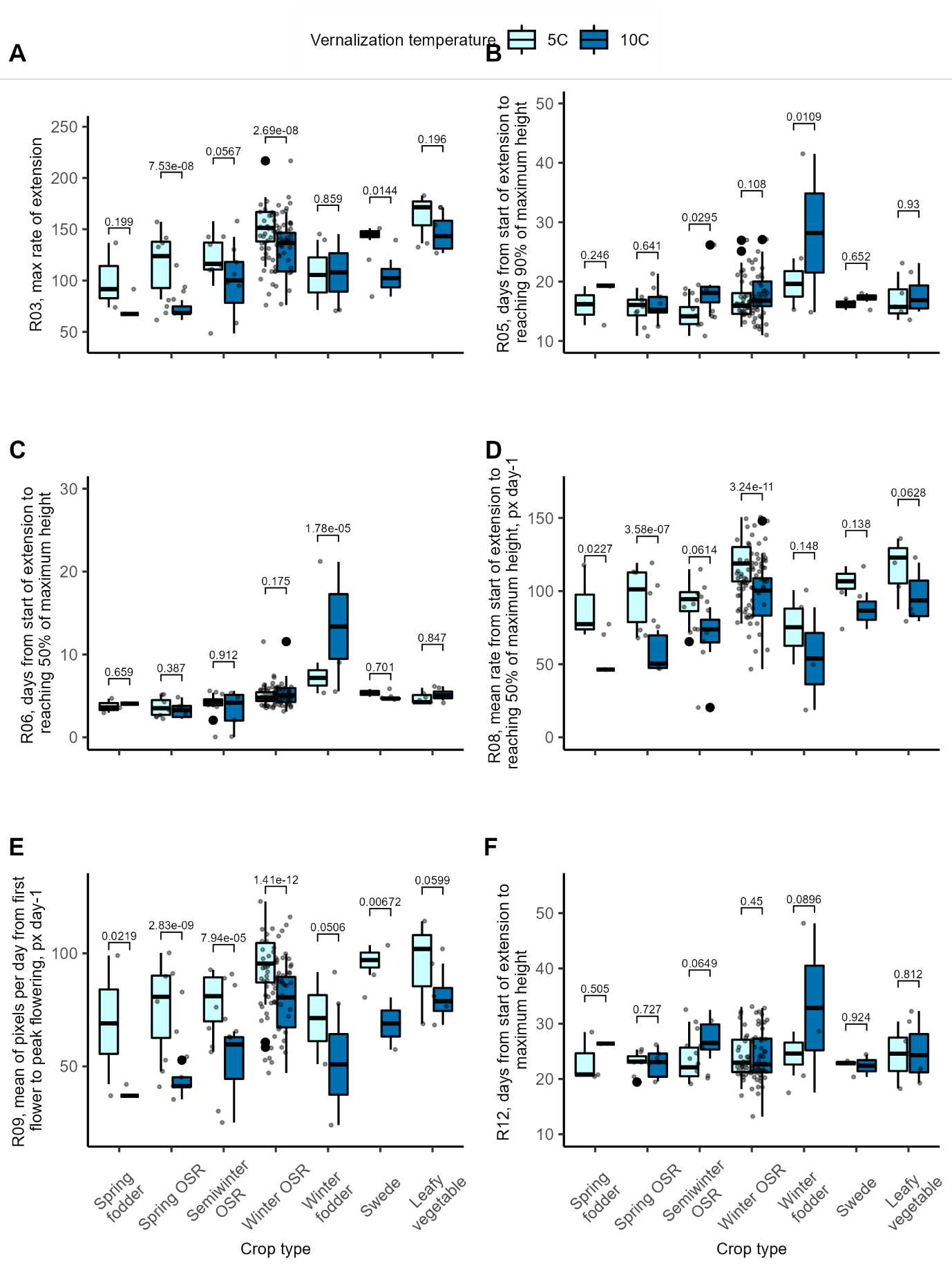

#### Supplemental Figure S7. Significant covariance between different traits within each genotype in response to different vernalization temperatures.

Covariation matrix showing relationships where p-value <0.05, with size and colour of marker indicating closeness and direction. A) traits after 5°C vernalization, B) after 10°C vernalization. C) trait results for 10°C minus results after 5°C vernalization, D) trait results for 10°C divided by results after 5°C vernalization. Aut = F10 (automated BBCH60 - ground truth BBCH51 measurement), G65 = G03 (ground truth BBCH60 - ground truth BBCH51), G51 = G01 (ground truth BBCH51), G60 = G02 (ground truth BBCH60).

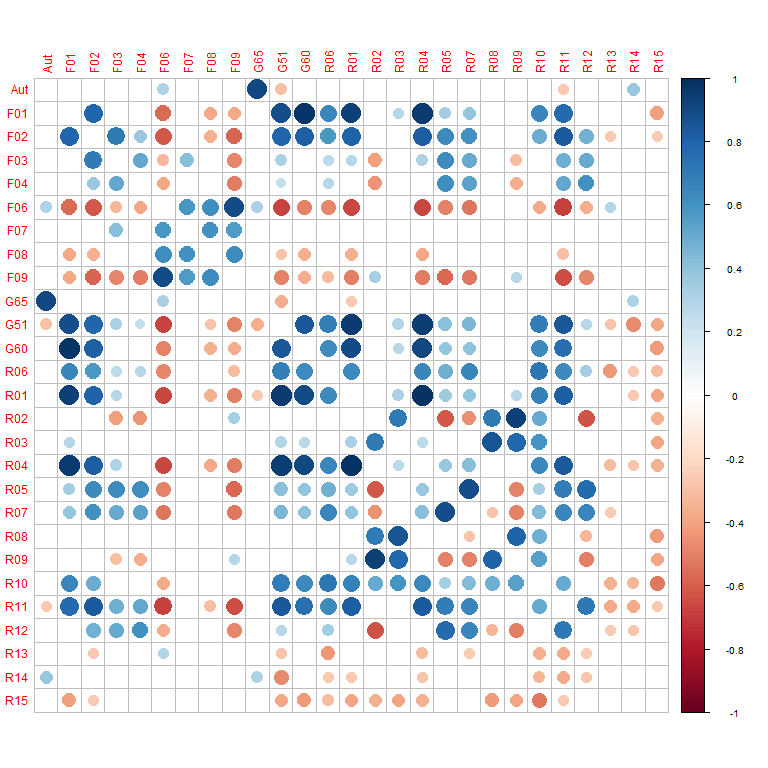

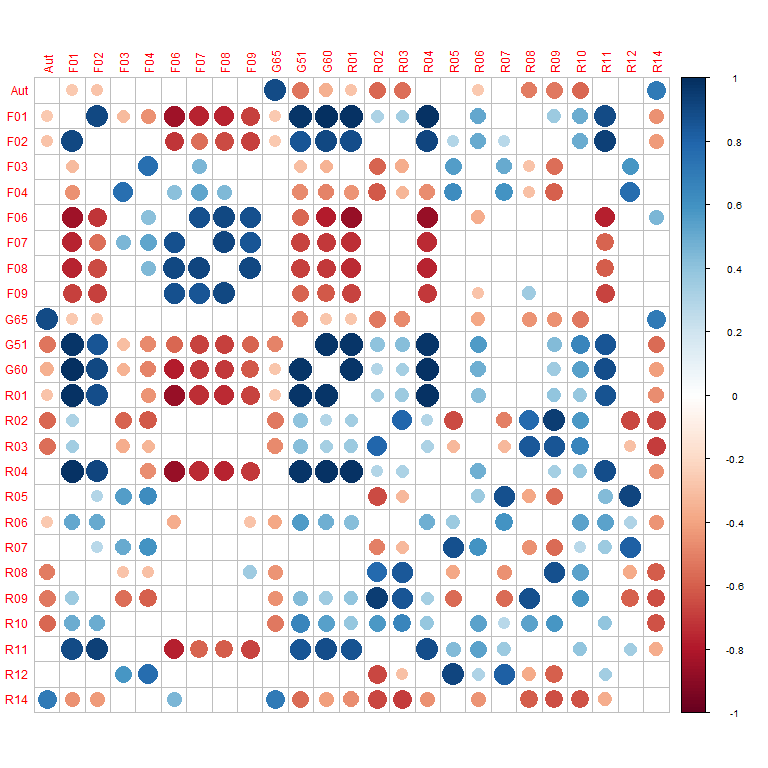

B

A

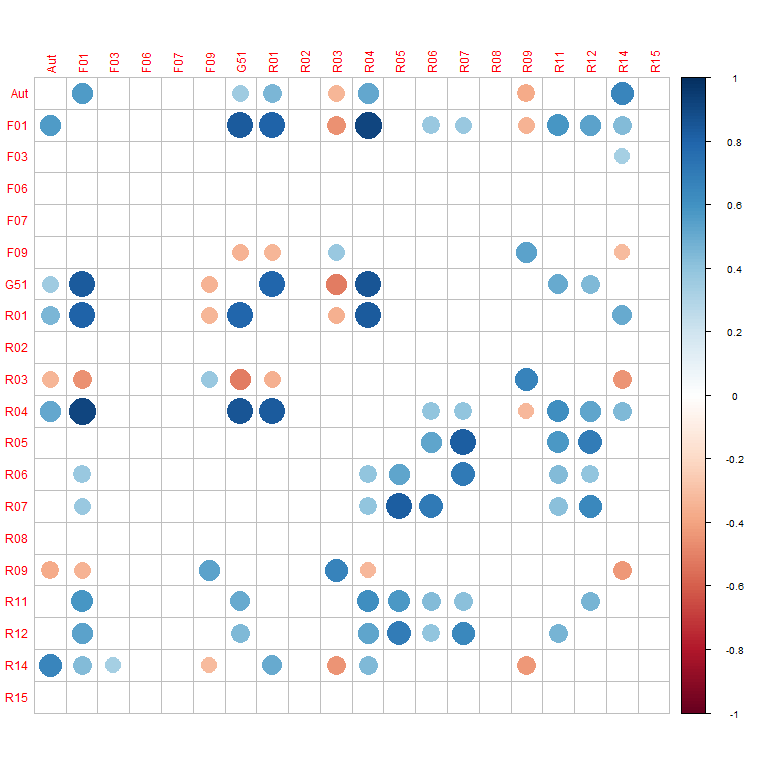

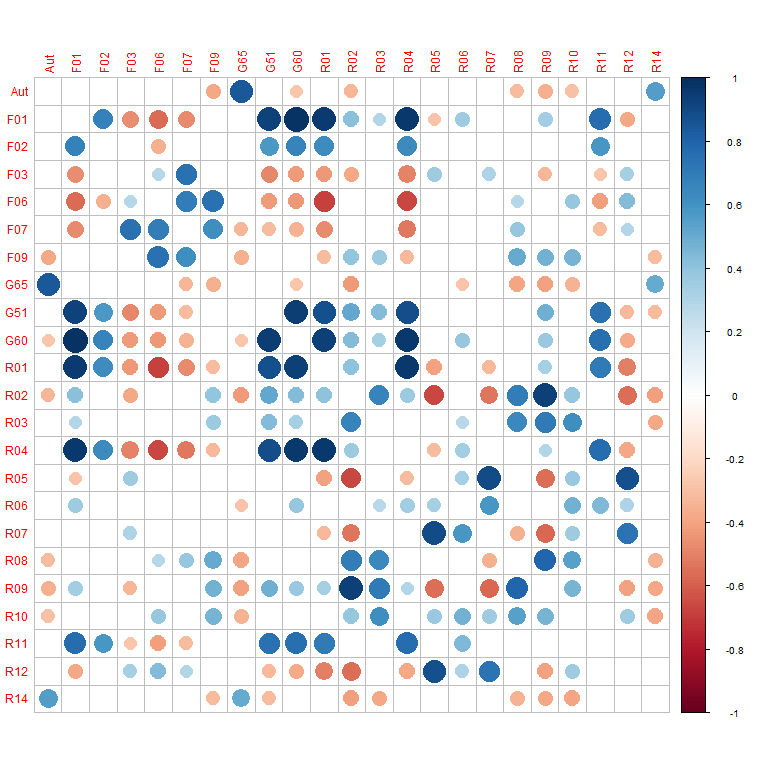

D

C

#### Supplemental Figure S8: Genetic variation at AGAMOUS-LIKE genes are associated with timing of peak flowering.

A) Presence/absence variation at *BnaAGL24.A01* is associated significantly with crop type, and with difference in time between buds visible and flower opening, as scored manually, after 10°C versus 5°C treatment (G03 at 10v5). (B) Haplotypes of *BnaAGL15.A03* are linked to the days to maximum height (R11) after 10°C treatment. Although five haplotypes were detected for *BnaAGL15.A03*, one was only represented by one accession and has been excluded. Points represent estimated marginal means per accession (see Methods). Violin plots showing mean (black circles) plus standard error (black lines), p values are FDR corrected post-hoc T tests. N varies per treatment and trait, but maxima are: Spring fodder = 3, spring OSR = 8, semiwinter OSR = 8, winter OSR = 42, winter fodder = 2, swede = 2, leafy vegetable = 6.

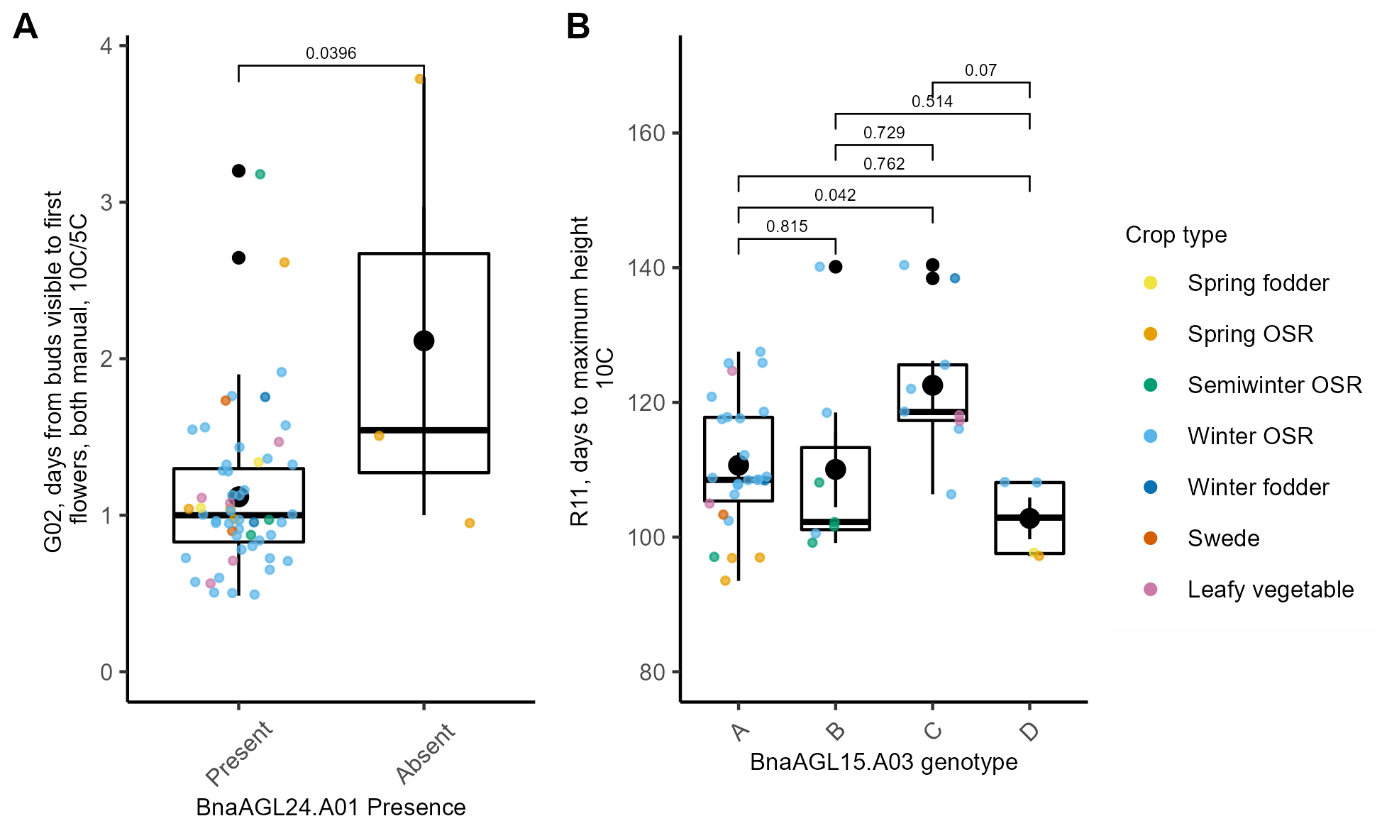

#### Supplemental Figure S9: Genetic variation at *BnaFLC.C02* is associated with timing of peak flowering.

Expression of *BnaFLC.C02* is linked to the peak flower timing trait after 10°C (p = 8.14 x10^-4^ for fixed effect of expression) (A) and in turn probably influenced by the *BnaFLC.C02* haplotype (p = 2.298 x10^-6^ of fixed effect of haplotype) (B), although linkage of haplotypes directly to the trait itself is not significant (p > 0.2 for fixed effect of haplotypes on F02 at 10°C) (C). Individual points are estimated marginal means per accession. A, line is regression. B, C, violin plots showing mean (black circles) plus standard error (black lines), p values are FDR corrected post-hoc T tests.

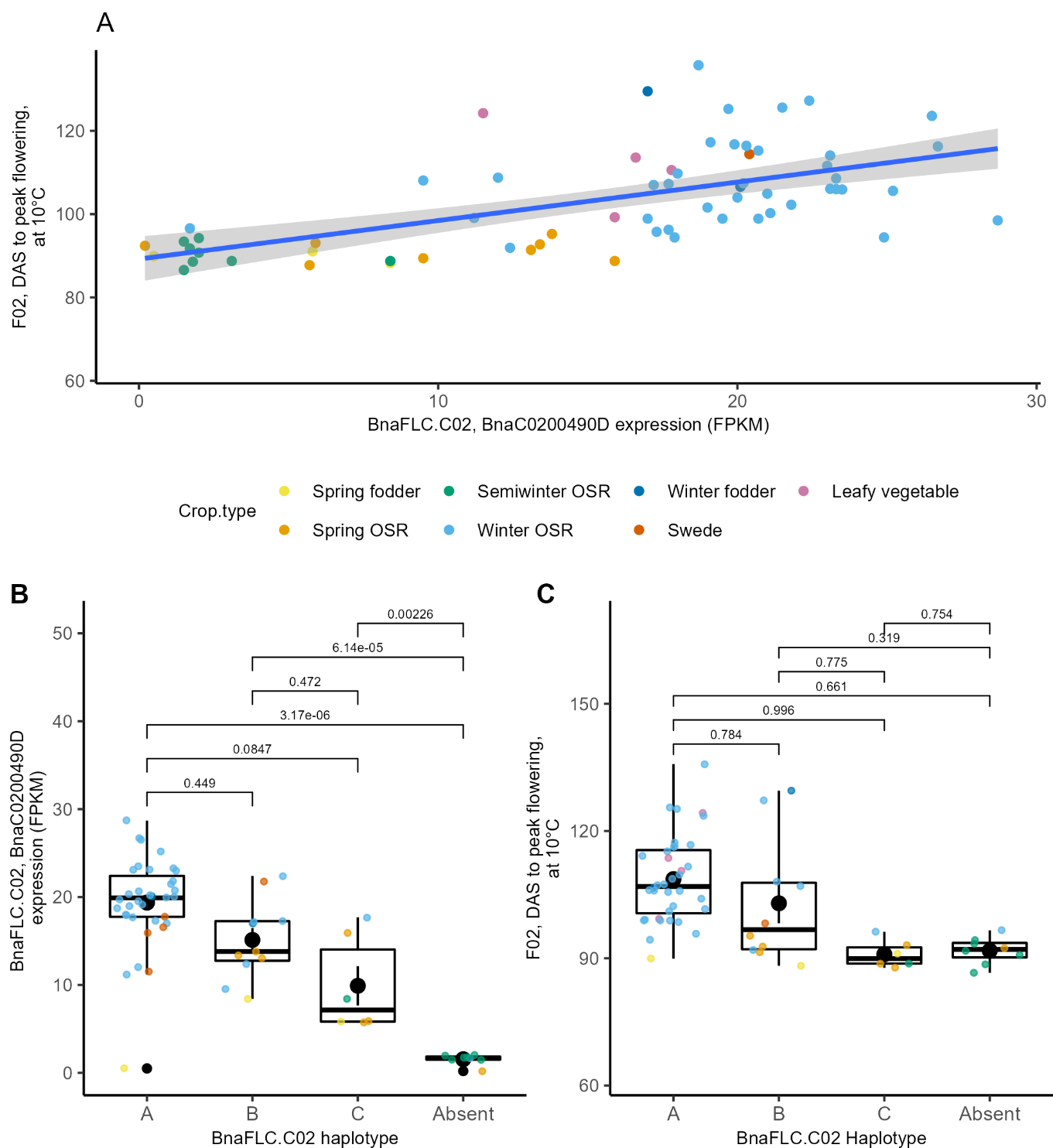

### Supplemental Tables

Supplemental Table S1. Genotype, treatment, glasshouse location and trait data used for analysis for all plants after 5°C vernalization.

Supplemental Table S2. Genotype, treatment, glasshouse location and trait data used for analysis for all plants after 10°C vernalization.

Supplemental Table S3. Genotype, glasshouse location and trait data calculated for 10-5 paired-plant comparison of temperature response.

Supplemental Table S4. Genotype, glasshouse location and trait data calculated for 10v5 paired-plant comparison of temperature response.

Supplemental Table S5. Significant SNP markers.

Supplemental Table S6. Significant GEMs.

Supplemental Table S7. Traits that do not present any significant results.

Supplemental Table S8. Significant eGWAS SNPs.

Supplemental Table S9. Genotype information.

Supplemental Table S10. Haplotypes, SNP calls and RPKM for Figures 6, S8, S9.

Supplemental Table S11. Haplotype and *BraBRAVE.A03* cDNA sequences.

Supplemental Table S12. Primers used.

### Supplemental Methods

#### Data Acquisition

The LemnaTec Scanalyzer occupies two glasshouse growing compartments (20 × 5 m) both containing seven parallel conveyor belts each holding up to 63 sample cars, giving a total of 882 cars. Images can be taken daily by transiting plants through a third compartment.

Whole plant side view and top view RGB images (cameras: Basler Pilot piA2400_17gc, 2056 × 2454, 5 MB; lens: Pentax TV Zoom, C6Z1218M3-5, 12.5 – 75 mm) are collected at intervals throughout the experiments. Image acquisition was initially daily but, as this became difficult to achieve with large plants, weekend imaging was suspended later in the growth cycle. Lighting in the RGB imaging cabin is provided by fluorescent tubes (OSRAM Lumilux Cool Day HE 28W/865; side view: 37 tubes; top view: 18 tubes). Uniformity of side view image lighting was slightly compromised in order to also capture top view images. Image exposure was optimised for detection of pixels related to flowering. Images were captured against a matt black background to reduce interference from chromatic aberration inherent in the standard LemnaTec design.

Images were saved in PNG format (RGBA; top view field of view 110 × 90 cm at height of pot; side view configuration set to a maximum plant height of 200 cm). LemnaTec configuration conditions are show in Supplemental Methods Table 1. Empty compost pots were included to provide a background image used in the initial segmentation (step 1).

For ground truth scoring of flowering time, plants were observed by eye up to three times a week and scored for visibility of buds at the apex (BBCH51; G01 in Table 1) or open flowers (BBCH60; G02).

#### Computational methods

##### Image analysis

Segmentation of High Throughput Phenotyping images and subsequent data analyses were conducted using bespoke software run using R versions x64 3.5.2 and 3.2.2 (R Core Team, 2018). Several R libraries were used to expedite this work:

‘rasterVIS’, version 0.45 (Lamigueiro, 2018)

‘png’, version 0.1-7

‘magick’, version 0.4 (using R version x64 3.2.2)

An initial sift removed those images where plants were not present, or where clear errors were noted during the experiment. The original software has been further developed into a pipeline, with possible application to other plant varieties.

##### Side view analysis of flowers

Initially, a single daily frame containing all treatments and replicates of each genotype was built from thumbnail images for each plant (400 × 495 pixels). Daily frames were compiled into videos to facilitate direct visual comparison of dynamic behaviours of each cohort.

Step 1

The flower segmentation algorithm starts with two side view images: the target plant, and an empty pot as a background. Red, green and blue channels of each image are processed independently, then recombined to make a mask where only yellowish pixels above certain thresholds are retained (0.50 and 0.75 are default limits). Two binary images are produced for each plant on each day: one where all yellowish pixels are included (∑pixel_>50_) which includes flowers and senesced leaves, and another where only brighter pixels are kept (∑pixel_>75_). The ratio ∑pixel_>50_:∑pixel_>75_ is used to determine whether original images contain senesced leaves: high ratios suggest more leaf material (Supplemental Methods Figure 1, Figure 2).

The masks are applied to the original image, and three colourised images are produced for every tenth date to assist with manual checking of the accuracy of segmentation (Figure 1).

Step 2

The periodic check images are stitched together for each plant. These combined images are inspected for minor adjustments needed to the colour channel parameters in step 1, to account for variation in the lighting

Step 3

Flowering pixel curves are plotted, including interpolated points when no daily image had been captured. Red dots represent the original yellow pixel count (flower + leaf); black circles are a minimum value of {∑pixel_>50_, 5×∑pixel_>75_} and green dots are interpolated results between black circles. Values for the green dots/black circles are used later in daily pixel counts (cases where leaves are present are relatively rare, but the approach described here acts as a good filter to remove such artefacts).

Flowering curves are parameterised to generate the nine digital phenotypes described earlier (Table 1, F01 to F09). The idealised curve (Figure 2A) shows the flowering features that are calculated within the constraints of first and peak flowering: F01, F03, F06, F07, F09. The calculations used are show in Supplemental Methods Table 2.

##### Top view analysis of flowers

Overhead images were used to evaluate flowering onset:

The first step uses a small section of the brassica rosette top view image (typically 400 × 400 pixels) and calculates binary masks in much the same way as for side view flowers (Supplemental Methods Figure 3). The area is kept small, and central to the rosette, due to possible interference from petals on the cabin floor. Further steps equivalent to the side view image analysis are conducted, but analysis was limited to flowering onset, as other metrics could not reasonably be calculated for these images:

(i) Flowers were at varying heights through the period, defeating the possibility of accurate area measurement (see Supplemental Methods Figure 4).
(ii) Older stems had a tendency to flop out of the ROI, and even out of the full field of view in some cases.

##### Side view analysis of raceme extension

An initial segmentation routine was developed to build a simple binary mask of each RGB plant image, with all background noise removed; this is used to calculate the height of each plant (Supplemental Methods Figure 5). By recording each height versus the full image timestamp (YYYYMMDDhhmmss) it is possible to plot the plant’s height versus time, and also its first order growth derivative (rate of raceme extension, [∆Height/∆Time]).

To avoid noise from the pre-extension period, the raceme extension is defined as having commenced two days before reaching 500 pixels tall (approx. 60 cm). This limit is based on observations of multiple growth curves across these experiments, was found to be successful in building strong growth rate curves for this panel, and readily identifies the features of date (MMDD) of growth onset and maximum growth rate.

Two curves are produced:

1. Daily height measurement in pixels.
2. Normalised growth rate in pixels per day (starting point: two days before the plant grows taller than 500 pixels).

The raceme growth curves are parametised to give the digital phenotypes R01 to R12, described in Table 1. Model growth curves and derivations for the raceme extension features are given in Supplemental Figure 3, with calculations used in Supplemental Methods Table 3.

##### Supplemental Methods Figure 1

Flow charts 1 & 2, flower segmentation using side view images

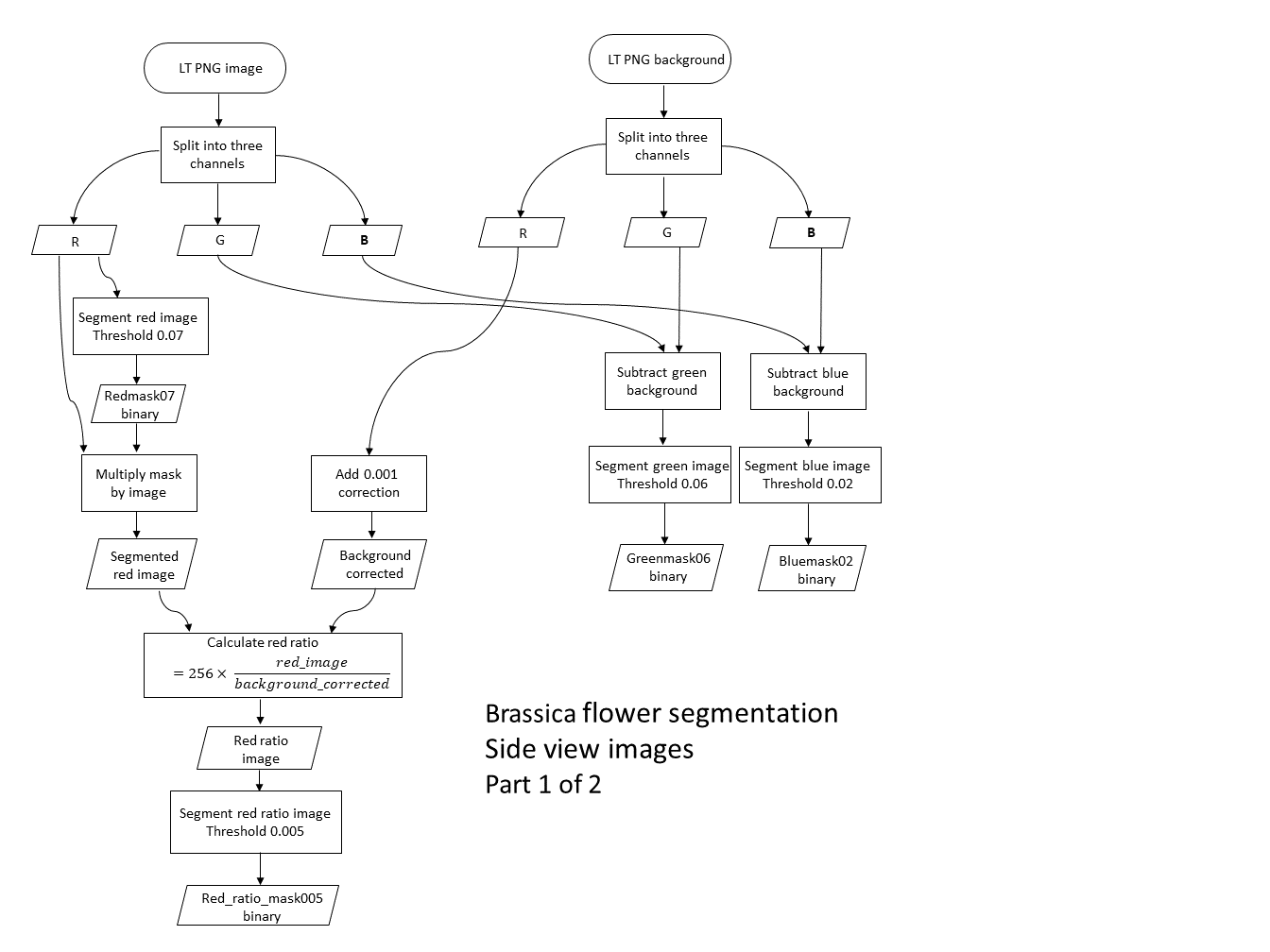

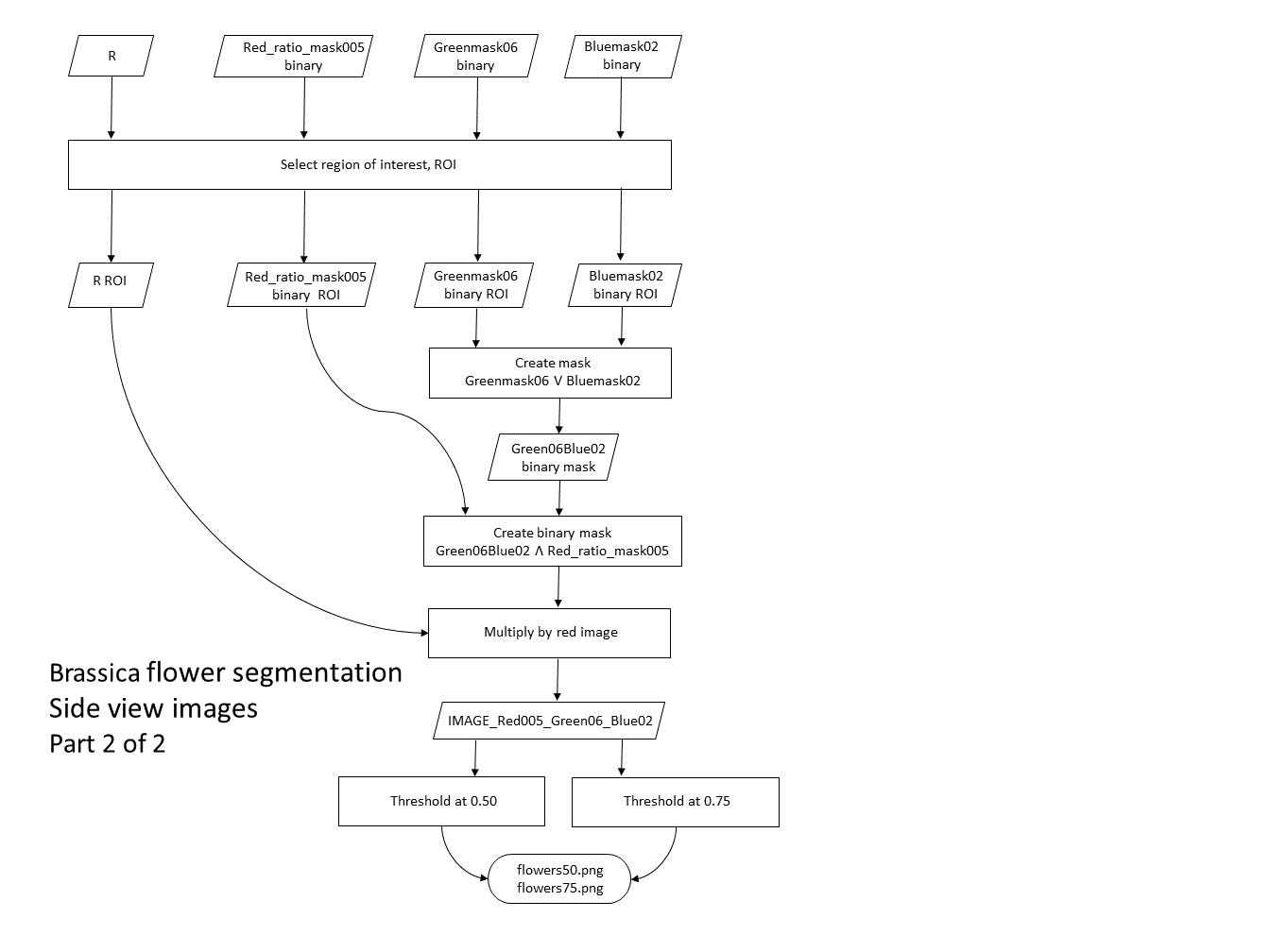

##### Supplemental Methods Figure 2

Isolation of flower (brighter) and leaf (darker) yellow pixels

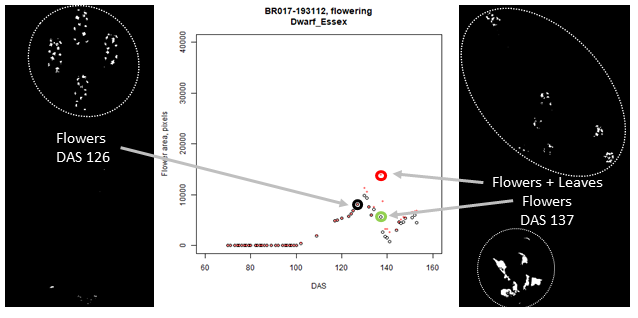

##### Supplemental Methods Figure 3

Flow charts 3, 4 & 5, flower segmentation using top view images

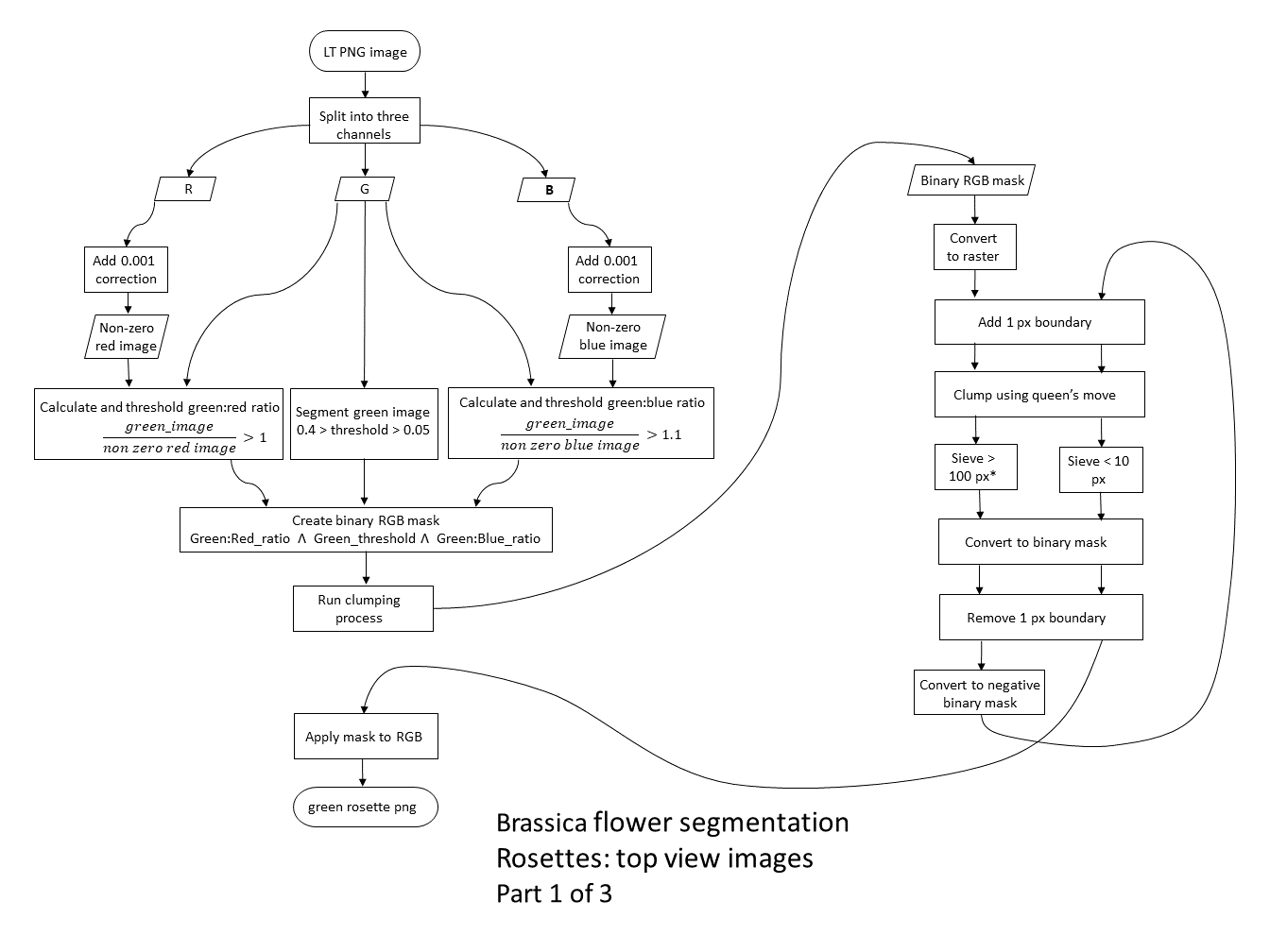

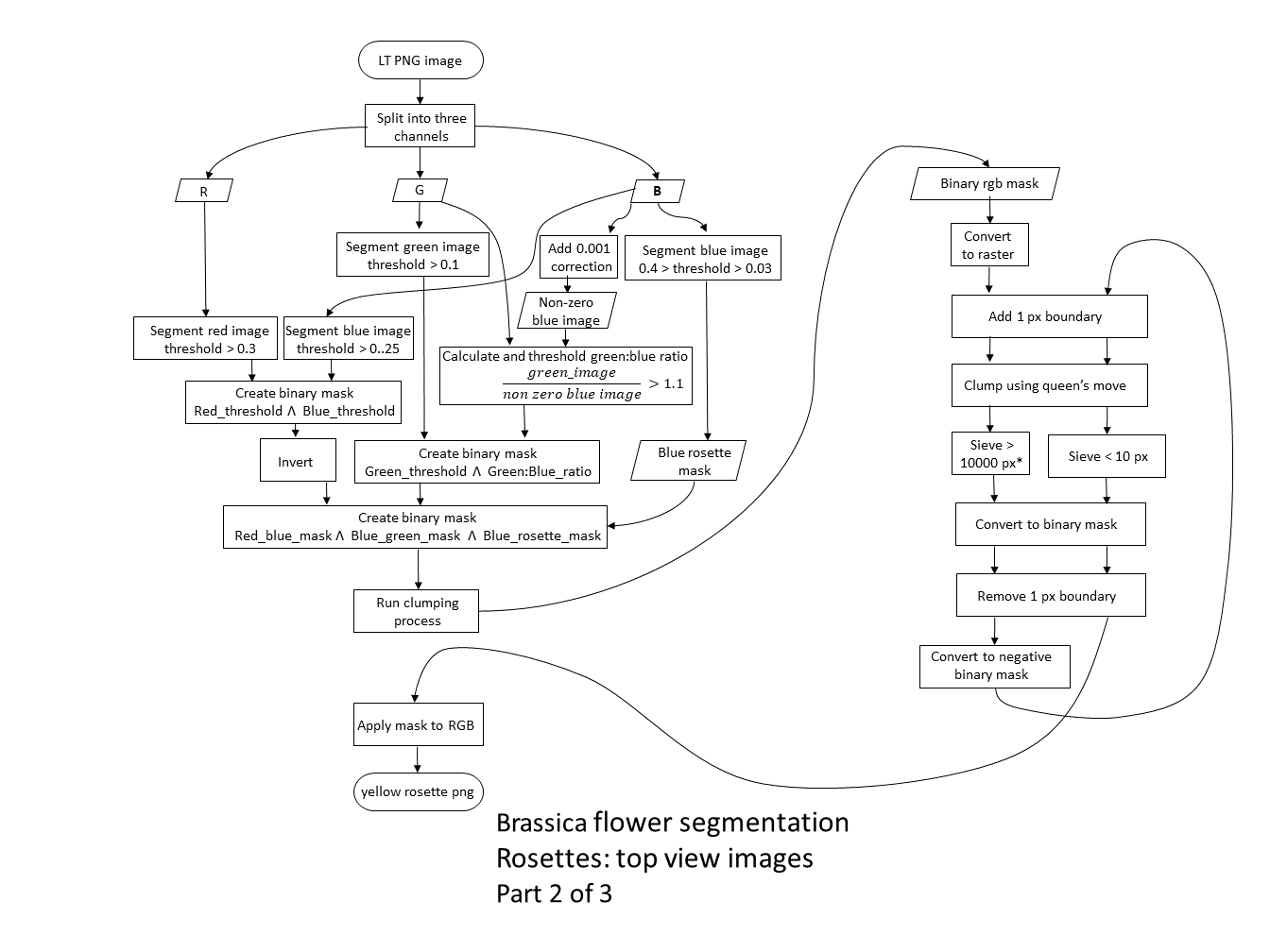

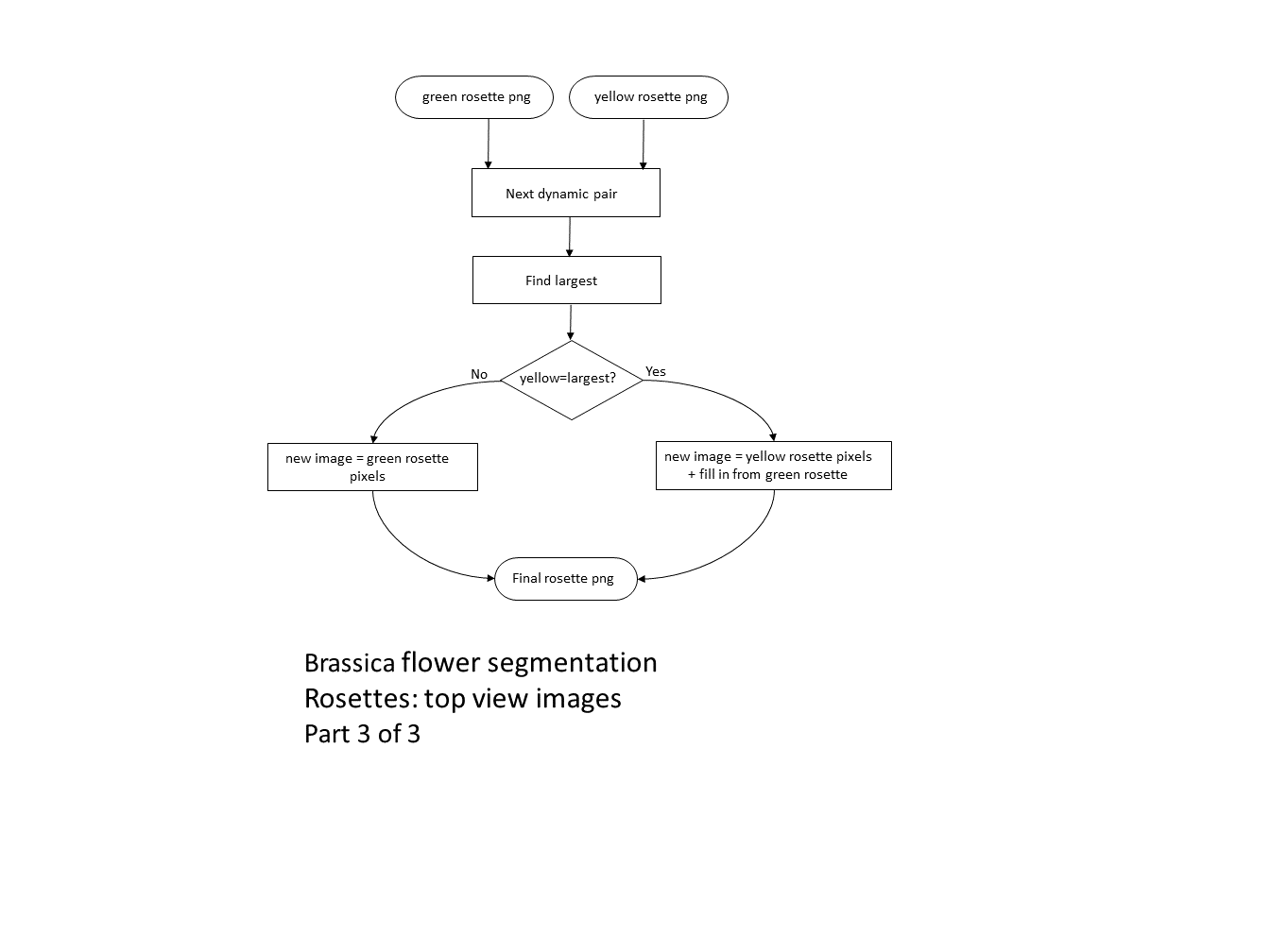

##### Supplemental Methods Figure 4

Variation of pixel density (relative pixel size) with height above the LemnaTec blue square pot holder, for top view images.

Variation of pixel density with height

| Distance above blue square, mm | Pixel density, px.mm-2 |
| --- | --- |
| 0 | 4.94618 |
| 81 | 5.1984 |
| 500 | 6.833 |
| 1000 | 10.0426 |

##### Supplemental Methods Figure 5

Flow chart 6, raceme segmentation using side view images

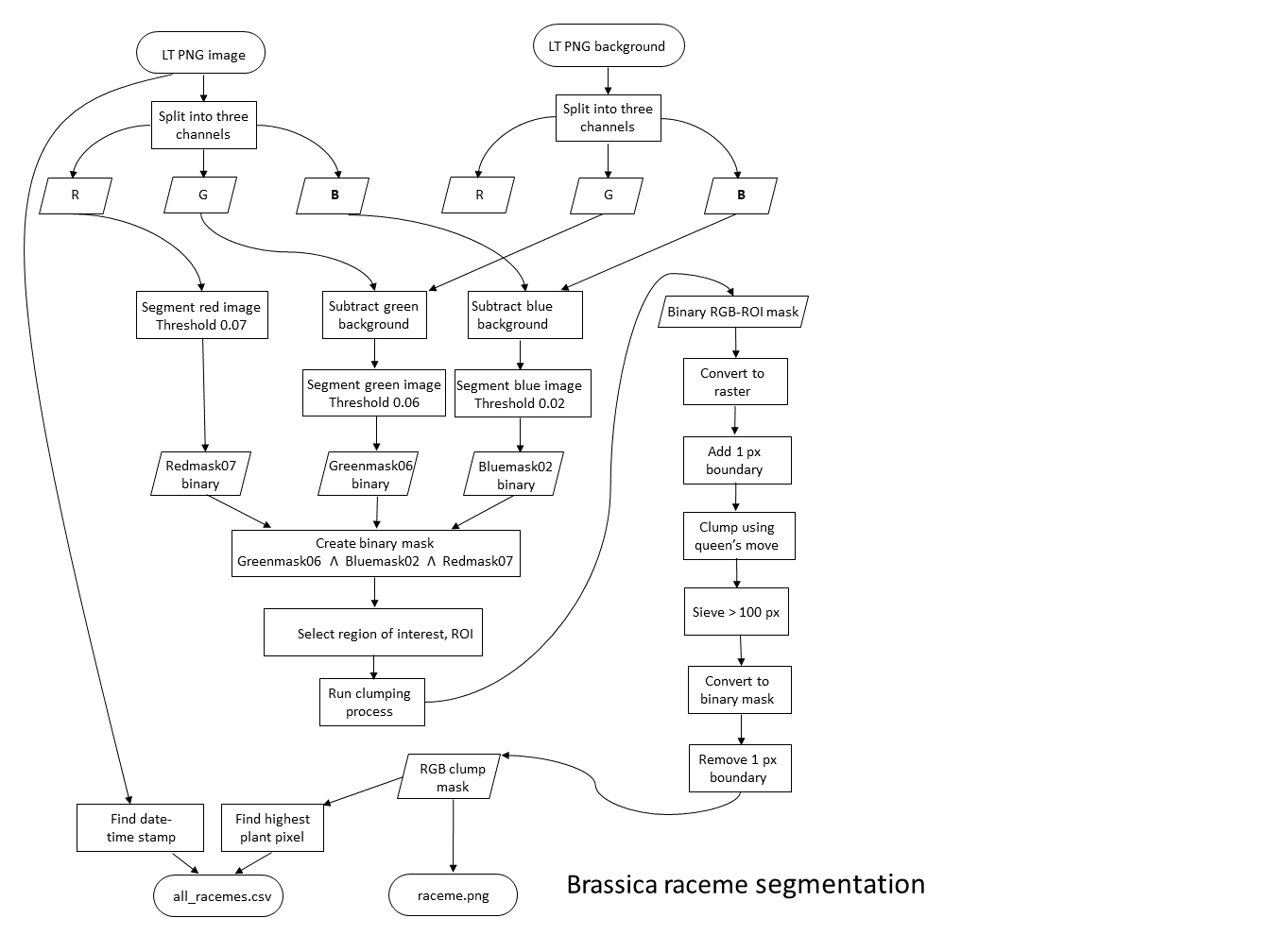

##### Supplemental Methods Table 1

LemnaTec configuration names and settings

| ID | Type | Angle, ° | Focus | Zoom | Exposure | Lifter height |
| --- | --- | --- | --- | --- | --- | --- |
| BRxxx_00_VIS_sv_000_A | Side | 0 | 4000 | 1500 | 500 | 1 |
| BRxxx_00_VIS_sv_045_A |  | 45 |  |  |  |  |
| BRxxx_00_VIS_sv_090_A |  | 90 |  |  |  |  |
| BRxxx_00_VIS_tv_000_A | Top | NA | 2850 | 2000 | 500 | 1 |

##### Supplemental Methods Table 2

| F01 | first_flowers_DAS |  |
| --- | --- | --- |
| F02 | peak_flowers_DAS |  |
| F03 | flowering_days_to_peak | *F02 – F01* |
| F04 | total_flowering_days | *F_end_ – F01* |
| F05 | flowering_finished |  |
| F06 | pixel.day_peak | $\sum_{F02}^{F02} {px}_{daily}$ |
| F07 | pixels.days_to_peak | $\sum_{F01}^{F02} {px}_{daily}$ |
| F08 | total_pixel.days | $\sum_{F01}^{F_{end}} {px}_{daily}$ |
| F09 | mean_pixels.day^-1^_to_peak | $\frac{F07}{F03}$ |

##### Supplemental Methods Table 3

| R01 | raceme_start |  |
| --- | --- | --- |
| R02 | raceme_extension_rate_90 | $\frac{{Px}_{90\%}}{t_{90\%}- R01}$ |
| R03 | max_extension_rate | $\frac{\Delta height}{\Delta timestamp}=max$ |
| R04 | max_rate_DAS |  |
| R05 | raceme_growth_period_90 | $t_{90\%}-R01$ |
| R06 | raceme_growth_period_50 | $t_{50\%}-R01$ |
| R07 | raceme_growth_period_75 | $t_{75\%}-R01$ |
| R08 | raceme_extension_rate_50 | $\frac{{Px}_{50\%}}{t_{50\%}- R01}$ |
| R09 | raceme_extension_rate_75 | $\frac{{Px}_{75\%}}{t_{75\%}- R01}$ |
| R10 | max_height |  |
| R11 | max_height_DAS |  |
| R12 | extension_days_to_max | $R11-R01$ |

##### Data analysis

The full set of growth parameters (G01 to G03, F01 to F10, and R01 to R15) are listed in Table 1 and Supplementary Tables S1-4. Ground truth parameters were compared to each other (G02-G01 = G03) and to the first flower data from the flowering curves (F01-G01 = F10)

For the flowering curves, features such as the dates of floral initiation and maximum flower-related pixels allowed ready calculation of parameters F01 to F03, and F06. The flowering period (F04) was not used for further comparative work, as some plants had to be removed from the platform before flowering had finished, as flagged by F05. When daily imaging jobs were missed, the flower-related pixel count for those days were linearly interpolated between known points; aggregation of these data allowed calculation of F07 and F09, with F08 also calculated for completeness, though not used for comparative work for the same reason as F04. A model flowering curve and parameter derivations are given in Figure 2A.

The raceme extension curves were initially plotted using only the available data, with no interpolated points. The mean daily growth rate was calculated for three interpolated periods, up to 50, 75 and 90% of each plant’s maximum height (R08, R09 and R02 respectively); the date and time of these occasions allowed the DAS to be calculated (R06, R07 and R05 respectively). The onset of raceme extension was defined as the interpolated timestamp 48 hours prior to the raceme passing through the 500 pixel height barrier. This definition was determined by observation, and was applied to both experiments. The maximum extension rate and date thereof were also derived (R03 and R04). Model curves and parameter derivations are given in Supplemental Figure 3.
